# The Hok bacterial toxin: diversity, toxicity, distribution and genomic localization

**DOI:** 10.1101/2025.08.07.668861

**Authors:** Andrés Escalera-Maurer, Adriana Messineo, Thibaud T. Renault, Elena Nicollin, Erika Castaneda-Sastre, Matthieu Brunot, Cléo Berrehail, Anaïs Le Rhun

## Abstract

The *hok/*Sok type I toxin-antitoxin system is encoded in chromosomes and plasmids of multiple Gram-negative bacteria. Current knowledge about *hok*/Sok regulation, toxicity and function originates mostly from the system present in the R1 plasmid, alongside a handful of homologs. The recent expansion of bacterial genome sequences calls for an updated analysis of *hok*/Sok diversity and distribution. Here, we used protein and DNA sequences as well as RNA structure to search for Hok homologs. Hok was detected in almost 80% of the available Enterobacteriaceae genomes and, more rarely, in three other bacterial families. Similarity, clustering analysis of thousands of Hok homologs unveiled high Hok sequence variability and uncovered clusters of novel homologs. Experimental validation of representative sequences revealed their toxicity, regardless of their localization. After scanning dedicated databases, we observed an enrichment of the *hok* sequences in prophages and large conjugative plasmids. Finally, we identified horizontal gene transfer events across genera, predominantly mediated by plasmids. Our findings yield a comprehensive and curated catalog of annotated sequences, detailing genomic localization, toxic activity and distribution across diverse bacterial genomes.

## INTRODUCTION

Bacterial toxin-antitoxin systems (TAs) are found across bacterial phyla (1). The toxin is a poisonous protein whose expression or action can be counteracted by the antitoxin encoded on the same genetic locus. Toxin production leads to the death or growth arrest of the cell encoding it. TAs’ exact biological function remains to be clearly defined. However, TA systems with characterized functions are involved in three processes: 1) maintenance of mobile genetic elements (MGE), such as plasmids or prophages by killing bacteria that have lost those elements during segregation – a mechanism named post-segregational killing (PSK) –, 2) defense against phages by killing the bacterial cell before phage propagation, and 3) formation of dormant cells (persisters) that can survive antibiotic treatment (2, 3). Eight TA types have been described with types I, II and III being the most extensively studied. In type I TAs (T1TAs), a small RNA antitoxin targets the toxin mRNA leading to mRNA degradation or translation inhibition. Type II TA antitoxins are proteins that impair toxin activity via a protein-protein interaction. Type III TAs consist of a small RNA antitoxin that inhibits a toxic protein (3, 4). Compared to type II and type III TAs, which are spread across several bacterial families, T1TAs appear to have a narrower distribution, potentially due to their small size and absence of recognizable domains, and are less frequently detected on MGE (1).

The most thoroughly investigated T1TA system is *hok*/Sok from the R1 plasmid, that became a paradigm for understanding post-transcriptional regulation by small RNAs (5). Hok (Host killing) is a 55 amino-acid toxic protein structured as a hydrophobic alpha helix, that forms pores in the bacterial inner membrane. Sok (Suppressor of killing) is an antitoxin RNA. A third gene within the locus, *mok* (Modulator of Killing), encodes a protein whose translation is coupled with that of *hok* and plays a key role in regulating its expression (6, 7). The chromosome of *E. coli* K-12 codes for five *hok* homologs: HokA, HokB, HokC (previously Gef), HokD and HokE. An additional layer of regulation was observed for the *hokB,* where its mRNA is adenosine-to-inosine (A-to-I) edited by the TadA enzyme, at codon 29, recoding a tyrosine into a cystein (8). Replacing codon 29 by a hard-encoded cystein codon, or overexpression of the TadA enzyme, drastically enhances HokB toxicity (8, 9). A similar phenomenon is observed in *hokC, hokD*, and *hokE* mRNAs for cystein 46. Indeed, a disulfide bond essential for HokB toxicity is formed between cysteines 29 and 46, explaining the link of RNA editing with toxicity (9, 10). In contrast, RNA editing is not required for toxicity of HokA, which already encodes both cysteins. Because *hokB* mRNA editing levels change in different growth phases, the requirement of RNA editing may act as an additional safeguard to prevent unwanted toxicity under specific condition (8).

The *hok*/sok system was originally discovered on the R1 plasmid as a mediator of plasmid maintenance through PSK (11, 12). In brief, the PSK mechanism proceeds as follows: when a daughter cell fails to inherit the R1 plasmid, rapid degradation of the Sok antitoxin leads to the activation and translation of the *hok* mRNA, which is more stable than Sok, causing Hok-induced cell death (13). Notably, treatment with rifampicin, an antibiotic that inhibits transcription, has also been shown to trigger Hok activation (6). Hok-induced PSK has been recently observed by live cell microscopy (14). It was also shown that overexpression of *hok*/Sok from a high copy plasmid protected bacteria against T4 phage infection (15). A role in persistence was observed for *hokB*, whose overexpression induces membrane permeabilization enhancing bacterial survival following antibiotic exposure (16). *hok*/Sok has also been proposed to increase survival under other types of stress in *Erwinia amylovora* (Peng et al. 2019). Finally, *hok*/Sok was also described as a plasmid competition system, as TA-encoding plasmids have an advantage compared to plasmids lacking TAs (17, 18). Regardless of its biological role, the killing activity of this system is of interest for potential biological and medical applications (19, 20).

Several searches for Hok homologs have been conducted previously. The first global identification of homologs was performed by Faridani *et al.* in 2006 using a TBLASTN approach (19). They found that Hok was confined to enteric and closely related bacteria, including bacterial pathogens, and showed that many bacteria harbored multiple copies. In 2010, Fozo *et al.* used customized and exhaustive TBLASTN and PSI-BLAST searches using the nucleotide sequence of the *hok* gene (21). Based on the search of T1TA homologs and the analysis of the Ldr/Fst and Ibs T1TA families, it was suggested that these systems are not spread by horizontal gene transfer (HGT) (21). In 2012, Steif and Meyer used RNA covariance models (CM) and identified 29 homologs of this system, demonstrating that taking advantage of the very stable and conserved structure of the *hok* toxin mRNA facilitates the identification of Hok loci (22). In 2017, Coray *et al.* used hidden Markov models (HMMs) on the amino-acid sequence to study the distribution of Hok family proteins, further highlighting the narrow distribution of T1TAs compared to other TA types (1). More recently a BLASTn search was used, but this time limiting the search to the genomes of plant pathogens (23). Stemming from the search done in 2010, the T1TAdb lists Hok homolog mRNA and peptide sequences annotated based on key determinants of the mRNA structure and genetic organization of the loci (24). Currently, 172 *hok*/Sok loci are annotated in this database. It was recently proposed that TAs, and Hok in particular, are good predictors of plasmid taxonomy (Bethke et al. 2024).

Ten *hok* homologs have been investigated in more detail. Hok, SrnB, Flm, PndA (from plasmid R16 and R483) are encoded on plasmids, while HokA, HokB, HokC, HokD (previously RelF), and HokE are encoded on the chromosome (25). In contrast to Hok from plasmids, the chromosomally-encoded loci *hokA* (strain *E. coli* C), *hokB* (strain K12), or *hokC* (strain ECOR24) could not mediate PSK and were not activated by rifampicin, leading the authors to suggest that chromosomal *hok*/Sok systems are inactive (26). Overexpression of the coding sequences (CDS) from all studied homologs showed that they were all capable of inducing cell lysis, except for HokB and PndA from plasmid R483 (25). Of note, all Hok CDS in this study were fused with several tags potentially affecting toxicity. Overexpression of HokB CDS alone was previously also shown to be toxic while the native locus from PndA from plasmid R483 was able to mediate PSK (26, 27).

Despite the wealth of information regarding *hok*/Sok regulation and toxicity, compiled up-to-date information about the localization, distribution and variability of *hok*/Sok homologs is missing. In this work, we identified 44248 homologs of Hok (1747 unique peptide sequences) in 6044 chromosomes, 10353 plasmids, 3852 prophages and 2501 phages. In addition, we sought to provide a detailed description of the diversity and prevalence of Hok across different species. We described their localization in chromosomes, plasmids and (pro)phages as well as their association with defense systems. We then tested experimentally the toxicity of diverse Hok representatives. Finally, we analyzed the occurrence of identical Hok sequences in distant bacteria, pointing to inter-genera transfer of Hok homologs via plasmids.

## RESULTS

### Prediction of Hok homologs shows high protein diversity

We first scanned the complete bacterial genomes of the RefSeq database for Hok homologs taking as input the Hok protein and mRNA sequences listed in the T1TAdb (24, 28). To increase the number of detected homologs we combined three methods using nucleotide (blastn), protein (MMseqs2) or mRNA structure and sequence (Infernal) (29–31). We reannotated all identified Hok CDS and extended each hit to include the closest start and stop codons, also annotating the Shine-Dalgarno (SD) sequence upstream the CDS when possible. We then removed degenerated sequences *ie.* CDS shorter than 40 amino-acids or containing internal stop codons (respectively accounting for only 5.1 and 2.6% of the unique CDS). We finally used CLANS to detect outlier sequences not connected to the main cluster (represented 0.43% of unique peptides) (32) (Fig. S1). CLANS generates a graph with nodes representing sequences and edges connecting sequences based on pairwise similarity scores. The distance between nodes are proportional to their pairwise similarity, allowing the visual identification of clusters and removal of outliers, which are likely false-positive hits. Closer inspection of these sequences revealed that the outliers were part of larger proteins with annotations unrelated to T1TAs, and therefore false-positive hits. The overall number of false-positive hits was very low.

We identified a total of 32,532 loci containing 1264 distinct peptides in the RefSeq database (Table S1). We were able to predict a putative SD sequence and a start codon for 84.5% of the peptides, while 10.2% contained only a start codon (without SD) and 5.3% did not contain either (Table S1). Approximately one third of all loci were identified by the three methods. The tool with the most diverse hits was Infernal, retrieving 85% of all hits and 89% of all unique peptide sequences (Fig. 1A). As Infernal uses both sequence and structure for prediction, we tested whether RNA structure was the key feature driving hit identification by Infernal, we performed a search that ignored the secondary structure. Only 8% of the sequences, corresponding to 16% of unique peptides (and 9 60% identity clusters), were missed when RNA structure was not considered, indicating that most of the hits could be found using RNA sequence alone. Therefore, although Infernal found most of the hits based on the sequence alone, mRNA structure contributed to increase sequence diversity. To get an overview of sequence diversity we built an all-against-all protein identity matrix. We observed a high variability of Hok sequences, with a mean percentage identity of 41.5%. Approximately 10% of the pairwise comparisons displayed an identity of less than 30% (Fig. 1B).

**Figure 1.**
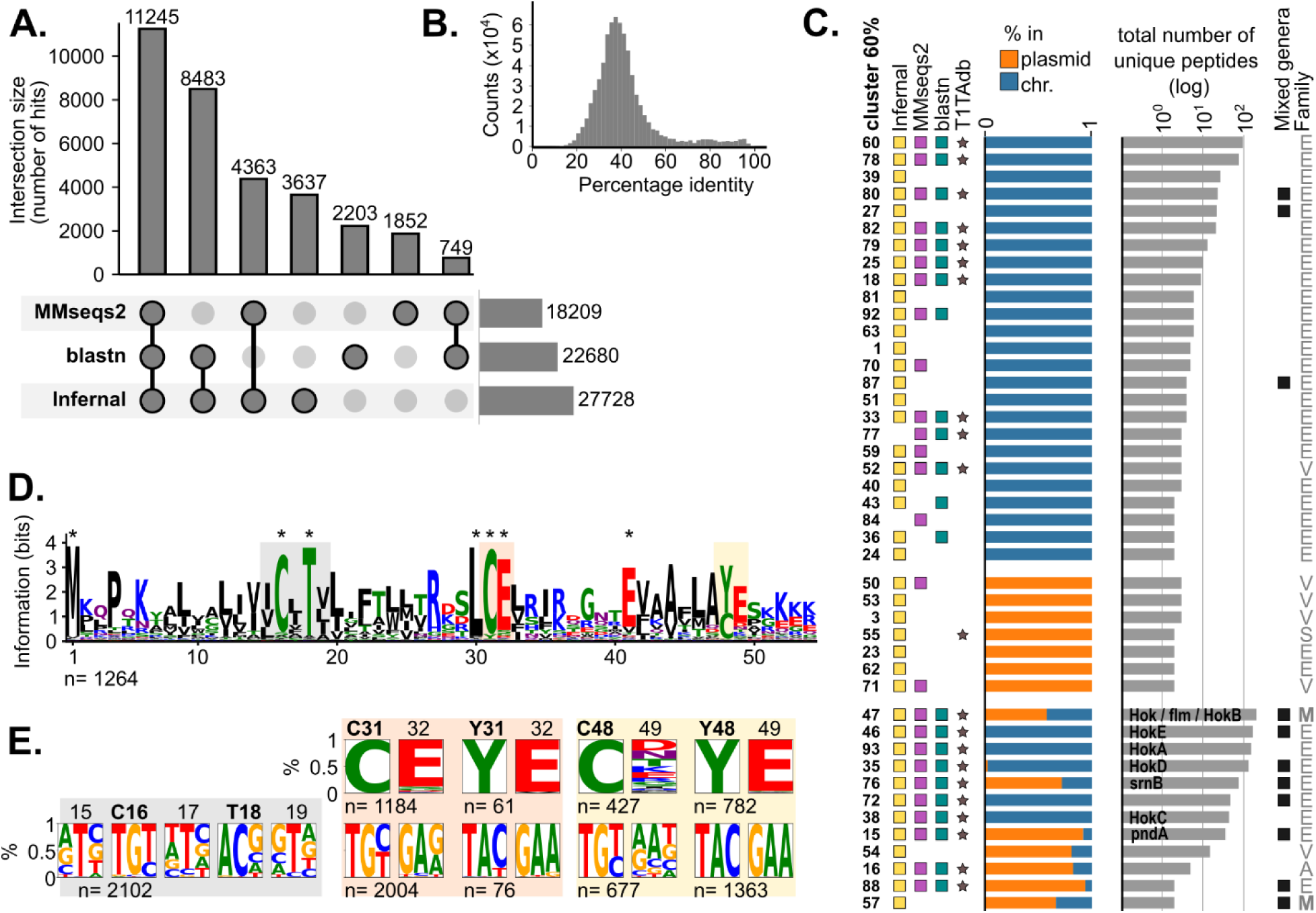
Identification and variability of *hok* homologs. **A.** UpSet plot summarizing the number of hits obtained by each of the three search methods against the complete bacterial genomes in the RefSeq database. The numbers above the bars indicate their corresponding values. Non-redundant refers to distinct peptide sequences. Hits numbers represent total of non-overlapping hits. **B.** Histogram of the distribution of Hok protein identity values in an all-against-all comparison matrix. **C.** Details on the 60% identity clusters of Hok containing more than one unique peptide. The symbols on the left indicate whether a peptide in that cluster is present in the T1TA database (T1TAdb, gray star), or was found by each of the methods: blastn, MMseqs2 or Infernal (green, purple and yellow squares, respectively) (see also Table S1). The bar plot on the middle shows the percentage of Hok loci from plasmid (orange) or chromosomes (blue). The total number of Hok unique peptides in each cluster is indicated by the bar plot on the right. The plot was visualized using iTOL. Previously experimentally studied homologs (HokA, B, C, D, E, Hok R1, *srnB, flm* and *pndA* R16 and R483) are indicated inside the bars corresponding to their cluster. On the right side, the clusters that contain sequences present in more than one genus are marked as “mixed genera” with a black square. The bacterial families represented in each cluster are marked: E (Enterobacteriaceae), V (Vibrionaceae), S (Shewanellaceae), A (Aeromonadaceae) and M (mixed, containing sequences from more than one family). **D.** Logo of the Hok amino-acid sequence using the 1264 unique sequences, after removing positions with >90% gaps. Aminoacids conserved in >80% of the peptides are indicated with an asterisk. **E.** Nucleotide percentage for C and Y codons at positions 31 and 48 together with the following codon. The codon logo for positions 15-19 is also shown. The number (n) of unique protein and unique DNA sequences used for constructing each logo is indicated.

Because Hok sequences were too short and variable to construct a meaningful phylogenetic tree, we clustered the Hok sequences with a 60% identity threshold and obtained 61 clusters (Table S1). Again, Infernal was the tool that identified the most diverse hits, covering 95% of the clusters. Interestingly, 9 out of the 59 clusters containing Infernal hits were not identified when the RNA structure was ignored for the search (Fig. S7). This suggests that these sequences were found based on RNA structure rather than sequence. The number of loci and unique peptide sequences varied greatly between clusters, and most sequences were concentrated in 16 clusters containing more than 10 unique sequences (Fig. S7). All experimentally characterized Hok systems belong to clusters with high number of sequences that are found both on plasmids and chromosomes (Fig. 1C). These include Hok homologs from plasmids R1 and R16, R483 (*pndA*), and F (*snrB*) as well as the HokA to E chromosomal homologs. The fact that we obtained so many clusters sharing less than 60% identity reflects the high diversity of Hok peptide sequences. The T1TAdb contains 82 distinct sequences distributed in 23 clusters, excluding inactive sequences. Although most of the clusters with more than 10 unique sequences had at least one representative in the T1TAdb, four of these clusters consisted entirely of newly discovered sequences (Fig. 1C). Thus, our search greatly expanded the quantity and diversity of known Hok sequences. Most of the peptide diversity was found on chromosomes. Indeed, 50% of the clusters, containing 74% of unique peptides, were composed of sequences not found in plasmids. In addition, 29% of the clusters contained sequences from both plasmids and chromosomes and 21% were only represented in plasmids. Only 35 unique sequences (7.3%) were found in both plasmids and chromosomes, suggesting that there is little transfer from chromosomes to plasmids and vice-versa, or that transferred sequences quickly accumulate mutations (Fig. S3A). Despite the tendency of plasmid and chromosome Hok sequences to cluster separately, no obvious differences set them apart at the amino-acid level (Fig. S3B, Files S1 and S2).

All of the clusters, except two, contained sequences present only in one family. Fifty clusters composed of only loci from Enterobacteriaceae, 6 from Vibrionaceae, one from Aeromonadaceae and Shewanellaceae (Fig. 1C). Similarly, most clusters are comprised of sequences from one genus. However, clusters with the most sequences tend to be spread across more than one genus (Fig. 1C). This suggests that sequences in large clusters could be transferred horizontally across genera.

To investigate the conservation of specific amino-acids, we aligned and created a logo of the 1264 unique peptides, after removing columns with more than 90% gaps from the alignment. Out of the 54 aligned amino-acids only 7 were conserved in >80% of the peptides (M1, C16, T18, L30, C31, E32 and E41) (Fig. 1D, File S3). At position 31 of the logo, a C is present in 92% of the sequences and a Y in 4.8%, the latter corresponds to *hokB* homologous sequences (Fig. 1D). The first nucleotide of the E32 codon is a G which, together with the Y31 UAC codon, forms the TadA recognition sequence (TACG) allowing RNA editing and Y-to-C recoding at position 31 (8). Codon 48 encodes a C in 34% and a Y in 62% of the peptides, respectively. When C is present, the following codon is not conserved. However, the E (GAA) codon is highly conserved when Y is encoded at position 48, once again allowing Y-to-C recoding by TadA RNA editing. (Fig. 1E). We also checked the codons encoding the conserved threonine T18 and did not find any codon conservation at this position, indicating that the protein sequence rather than the DNA/RNA sequence was important.

### Experimental validation of Hok peptides toxicity

The search for Hok homologs returned 1264 unique sequences that are potentially active. Indeed, we were unable to predict whether the peptides are toxic based on their sequence alone due to the lack of conserved domains in Hok. To examine the relationship between amino-acid sequence and activity, we first tested the toxicity of mutants of Hok R1 CDS. The toxicity of the CDS of conserved amino-acids mutated to alanine (namely C16A, T18A, L30A, C31A, E32A, E41A and E49A) was tested by assessing their ability to inhibit the growth of *E. coli* upon overexpression. The CDS sequences, spanning from the start to the stop codons, were cloned under an arabinose-inducible promoter while keeping the SD constant. The absence of conserved residues did not prevent toxicity, unlike a construct carrying a truncated Hok (R1*Stop) (Fig. 2).

**Figure 2.**
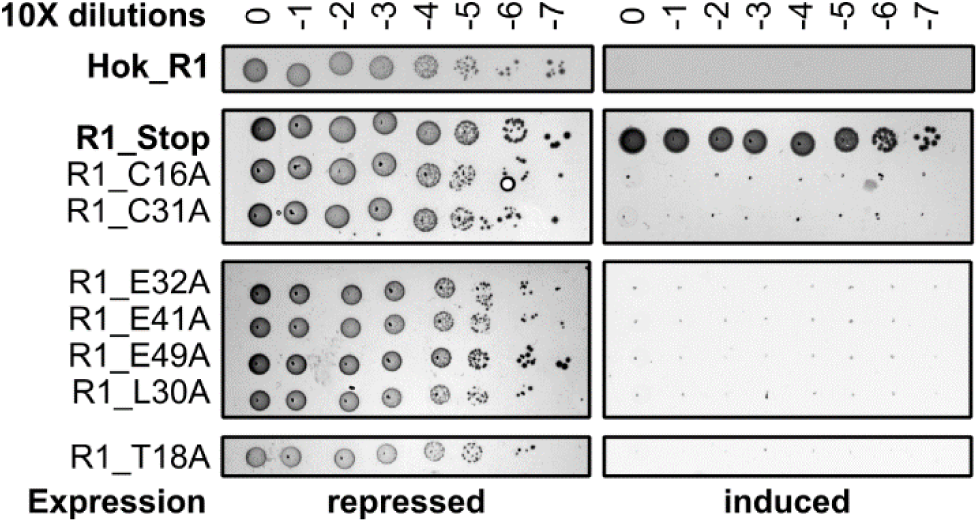
Toxicity of Hok mutants. Drop assays of *E. coli* carrying the control vectors (Hok R1 or Hok R1 containing a stop codon) or a plasmid encoding alanine-substituted Hok R1 mutants, under repression with 0.2% glucose (left) or induction with 0.01% arabinose (right) of *hok* expression. Drops of undiluted bacteria and 10-fold dilution until 10^-7^ were plated. No growth indicates toxicity.

We then selected representative Hok homologs to test their toxicity experimentally. Those homologs were selected from the 16 most abundant clusters (containing more than 10 sequences), and from 14 clusters containing only one Hok sequence (Table S2). Despite repeated efforts for cloning cluster 39, we only obtained mutated sequences, suggesting that this homolog is toxic. To validate our assay, we used sequences that had been previously characterized in the literature as controls. These included: Hok from plasmid R1 (47_377), HokA (93_1378), HokC (38_529), HokD (35_697), HokE (46_966) and HokB (47_188), PndA from plasmid R483 (15_1414) and Flm (47_385) shown to exert a toxic phenotype (26, 27, 33–36). Out of the 45 homologs tested, 42 were toxic (Fig. S4). Only sequence 19_179, 60_1482 and 69_1540 were not toxic. The reason for the lack of toxicity of those sequences was not obvious. For example, sequence 60_1235 from the same cluster than 60_1482 was toxic. The toxicity was also independent of the number of sequences in the cluster indicating that clusters with one sequence do not consist of degenerated/inactive Hok. The localization of the selected Hok in the chromosome or mobile genetic elements did not affect the toxicity in our conditions. This means that that most identified Hok proteins are potentially functional (Fig. S5).

### Hok prevalence is highly variable between Enterobacteriaceae genera

Previous bioinformatics studies have investigated the distribution of *hok*/Sok systems in bacterial genomes (1, 21). However, detailed analyses of their abundance and localization had yet to be conducted. In accordance with previous reports, the great majority of Hok homologs was found in the Enterobacteriaceae family, with 75% of the chromosomes and 29% of the plasmids of this family containing at least one homolog (Fig. 3A). While a few examples were also detected in Vibrionaceae, Aeromonadaceae, Shewanellaceae and Pseudomonadaceae (Fig. S6A and B, File S4) the prevalence was between 10 and 13% for plasmids and less than 4% for chromosomes in those families (*hok* was not identified in the chromosome of Shewanellaceae) (Fig. 3A). One instance was found in a plasmid assigned to the Pseudomonadaceae family but was disregarded because a blast search returned 100% similarity to plasmids in Klebsiella and we did not detect any similarity to other plasmids from Pseudomonadaceae. We did not find any distinctive features in the amino-acid sequence of the Hok homologs outside Enterobacteriaceae (Fig. S6A and B).

**Figure 3.**
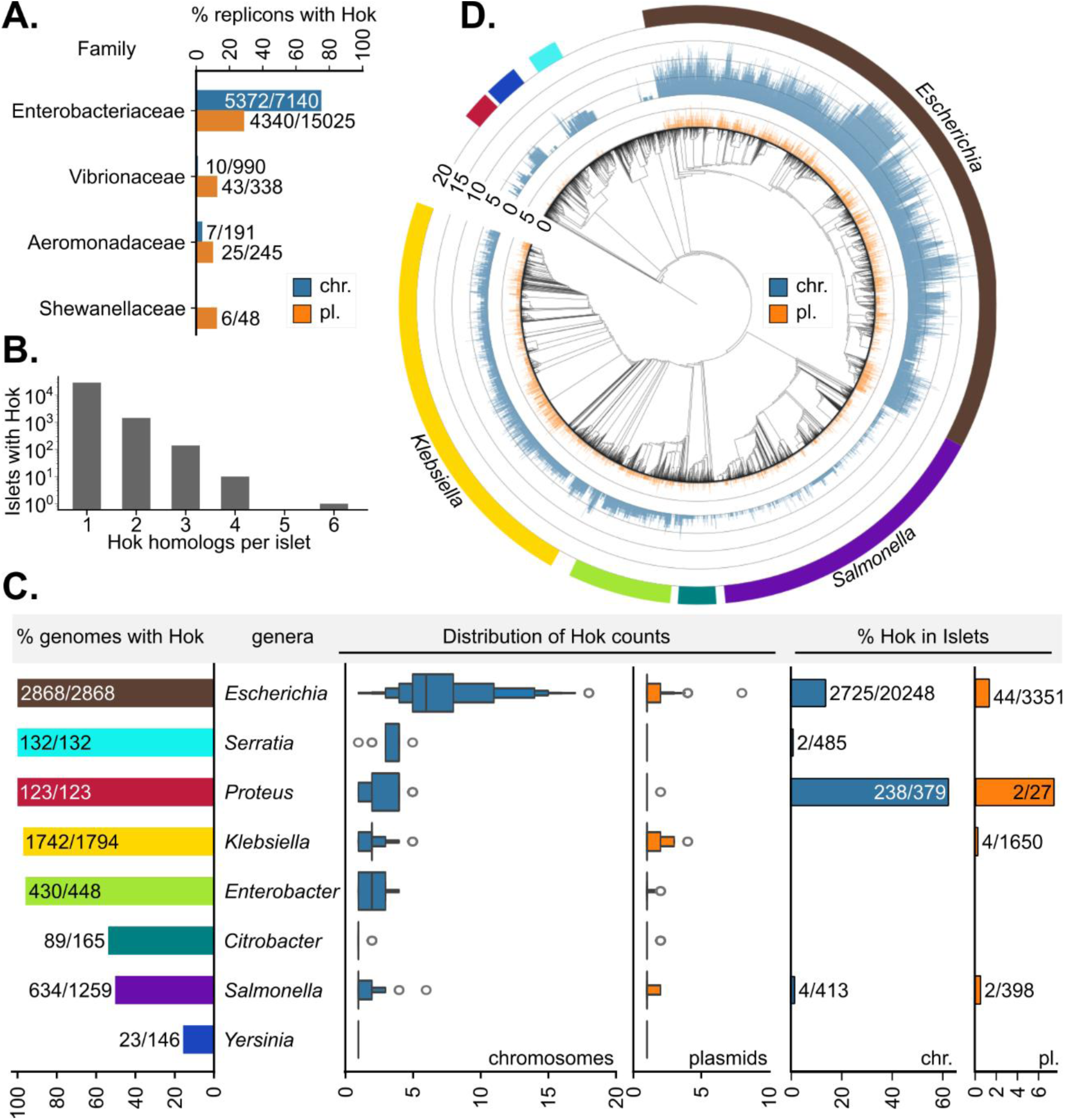
Distribution of *hok* homologs. **A.** Percentage of plasmids (orange) and chromosomes (blue) containing *hok* per bacterial family, the values on the bars are the number of *hok*-containing plasmids or chromosomes over the total number of searched plasmids or chromosomes, respectively. **B.** Number of islets containing *hok* genes within 2 kb of each other. The x-axis indicates the number of homologs contained in each islet. **C.** Left: percentage of genomes with *hok* in Enterobacteriaceae genera (containing > 100 genomes), the number of genomes with *hok* over the total number of genomes per genus are indicated on top of the bars. Middle: boxen plot indicating the distribution of the number *hok* loci in plasmids (orange) and chromosomes (blue) per genus. Right: percentage of *hok* homologs in islets (within 2 kb from another homolog), values are the number of homologs in islets over the total number of homologs. **D.** Phylogenetic tree of Enterobacteriaceae chromosomes where the search was conducted, constructed with GTDB and visualized with iTOL. Bars radiating from each tree leaf indicate the number of *hok* loci encoded in each chromosome (blue) or its associated plasmids (orange). The eight most abundant genera are indicated by the same colors used in C.

It was previously reported that T1TAs can be repeated multiple times in the same locus (21). Here we found that 10% of Hok were found in islets of 2 to 6 homologs each one less than 2 kb apart from another (Fig. 3B). Islets seem to arise by duplication of *hok* genes in the same locus rather than multiple independent insertions of distant *hok* homologs. Indeed, 98% of all islets contain peptides belonging to only one 60% identity cluster. However, in 96.3% of the cases, the replicons where the islets were located encode a Hok from a different cluster outside the islet, indicating divergent homologs are mostly located more than 2 kb apart.

The number of available genomes for most of the Enterobacteriaceae genera was limited. Therefore, we focused on genera with more than 100 sequenced genomes. The number of *hok* genes in chromosomal islets was less than 1% for all genera except *Escherichia* and *Proteus* where it was 13.5 and 62.8%, respectively (Fig. 3C). A similar pattern was observed for plasmids, albeit with 1.3 and 7.4% of *hok* genes found in plasmidic islets in these genera, respectively (Fig. 3C). Prevalence of Hok in these abundant genera also varied widely. For *Escherichia*, *Serratia* and *Proteus*, 100% of the analyzed genomes contained at least one *hok* copy (in the chromosome or one of its associated plasmids). The percentage of genomes with *hok* was also high for *Enterobacter* and *Klebsiella* with more than 95% of genomes encoding *hok*. In contrast, only half of genomes in *Citrobacter* and *Salmonella* contained a *hok* homolog (Fig. 3C). The genus with the lowest *hok* prevalence was *Yersinia* with only 15.8% of genomes containing *hok*. The median number of homologs per plasmid was one for all genera. The maximum number of homologs was in *Escherichia coli*, which harbored 8 homologs in a >381 kb plasmid, and up to 18 in the chromosome (median of 6) (Fig. 3C). The chromosome with the highest number of homologs belongs to *E. coli* O157:H7 (EHEC). The rest of the genera had a maximum of 2 to 6 homologs with a median of 1 per chromosome. To further investigate the distribution of homologs, we mapped them on the phylogenetic tree of Enterobacteriaceae genomes and observed that their distribution varies considerably across genera (Fig. 3D).

### *hok* gene is not enriched in the vicinity of defense systems

*hok*/Sok was shown to play a role in defense against phages (15). Because defense systems are known to cluster near other defense systems (37–39), we investigated whether Hok was preferentially found in proximity to known defense systems. For that, we first predicted defense systems in genomes with *hok* using Defense finder (40, 41). We then calculated the percentage of homologs encoded less than 20 kb from a predicted defense system. As a control, we computed the percentage of homologs that would be near defense systems if their distribution in each chromosome was random, keeping their numbers per chromosome unchanged. The percentage of *hok* homologs near defense systems was comparable to the expected value if their location was random (12 vs 9.4%) (Fig. 4). As an additional comparison, we repeated the analysis for the SymE/SymR T1TA, which is enriched near defense systems (42). Randomly distributing SymE homologs resulted in 5.6% being near defense systems compared to 96% when calculated using the actual coordinates (Fig. 4).

**Figure 4.**
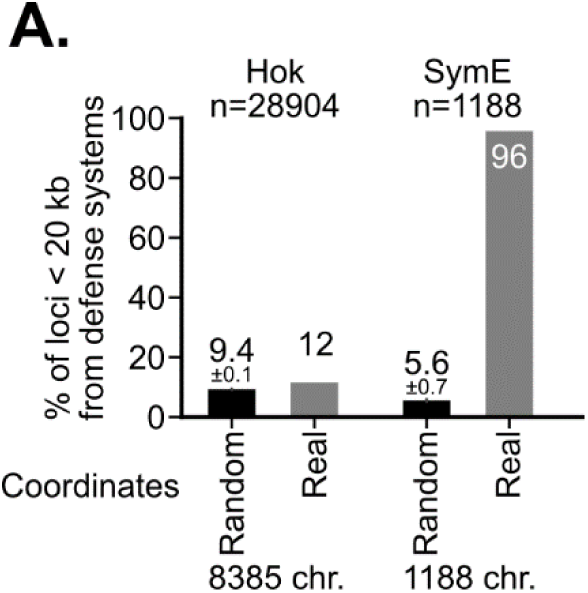
*hok* genes are not enriched next to defense systems. The percentage of loci located less than 20 kb from known defense systems found by DefenseFinder is displayed in the plot. “Random” indicates the percentage of *hok* homologs located near defense systems after randomizing their location, while “real” indicates the actual proportion of homologs near defense systems. For comparison, the real and randomized proportions of the SymE type I toxins adjacent to defense systems are shown. The number of homologs found in genomes with defense systems is indicated (n). Only chromosomes containing both TA (Hok or SymE) and defense systems were considered.

### *hok* genes are enriched in prophages

T1TAs are also involved in maintaining prophages in the chromosome (43). In addition, some anti-phage systems are encoded in phages (44). Therefore, we sought to investigate the presence of *hok* in phages. For that, we searched for homologs in the IMG_VR database, which contains a collection of viruses identified in publicly available microbial sequences, using the CDS predicted from the RefSeq searches (45). We found a total of 6290 Hok homologs (including 385 unique sequences) in phages and prophages (Table S1 and Fig. S7A and B). We also identified 9 new 60% identity clusters in this search. The great majority of Hok containing phages (96.3%) encoded one homolog and only 3.5% and 0.2% encoded 2 or 3, respectively. All *hok*-encoding phages belong to the Caudoviricetes class and infect Enterobacteriaceae almost exclusively (except for three phages infecting Vibrionaceae). Of all Enterobacteriaceae-infecting phages, 5.25% encode at least one Hok. That percentage ranges from 9.3% for the *Escherichia* genus to less than 0.1% for *Klebsiella* (Fig. 5A), which contrasts with the observed prevalence of 97% for *Klebsiella* genomes in the RefSeq (Fig. 3C).

**Figure 5.**
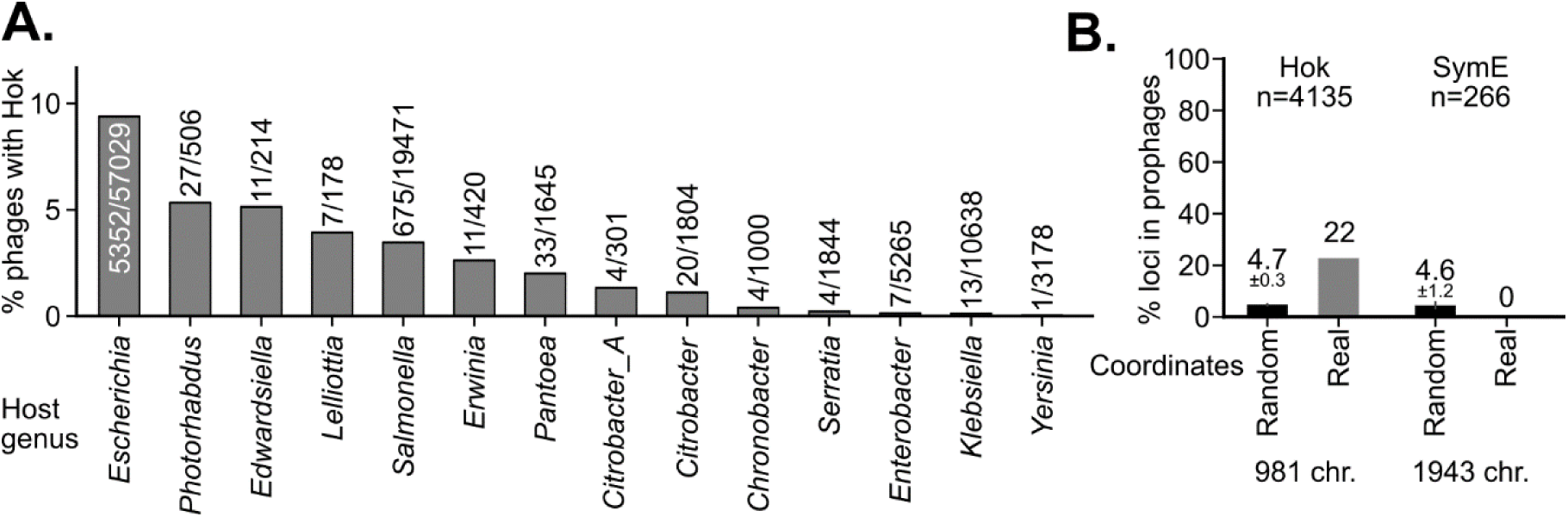
*hok* genes are enriched in prophages. **A.** Percentage of phages encoding Hok per bacterial host genus. The number of phages that code for Hok over the total number of phages are indicated on top of each bar. **B.** Bars represent the percentage of *hok* homologs in prophages (values are shown on top of each bar). “Random” indicates the percentage of Hok homologs located in prophages after randomizing their location, while “real” is the actual proportion of homologs in prophages. SymE type I toxins are not present in prophages. Only chromosomes containing both TA (Hok or SymE) and prophages were considered. Number of homologs (n), and number of chromosomes (chr.) are shown.

To test if Hok was preferentially found in prophages, we mapped all confidently-predicted phages on the chromosomes containing Hok. We found prophages in 16% of the 981 *hok*-containing chromosomes. In these, 23% of the 4135 Hok homologs were encoded in prophages, four times more than the 5% expected if *hok* homologs were inserted randomly in the chromosome (Fig. 5B). In contrast, SymE was not enriched in chromosomes containing both SymE and prophages, the expected percentage was 4.6% for the randomized SymE coordinates (Fig. 5B). Overall, this suggests that *hok* genes are enriched in prophages.

### *hok* loci are enriched in large plasmids

To perform a more detailed analysis of the distribution and prevalence of Hok in plasmids, we looked for Hok homologs in the IMG_PR plasmid database, containing homogeneously annotated plasmids (46) (Table S1 and Fig. S7A and B). Hok was found in plasmids associated with the same families than in the RefSeq search. The family with the highest percentage of *hok*-containing plasmids was Vibrionaceae with 18% (44/240) (Fig. 6A). The highest number of plasmids with Hok was detected in the Enterobacteriaceae family, though only 11.3 % of all plasmids contained Hok (Fig. 6A). In Enterobacteriaceae genera with at least 10 homologs, the percentage of plasmids with *hok* ranged from 32% (15/46) in *Proteus* to 8.5% (86/1008) in *Salmonella* (Fig. 6B). We identified 9 new 60% identity clusters in this search, reaching 79 clusters in total.

**Figure 6.**
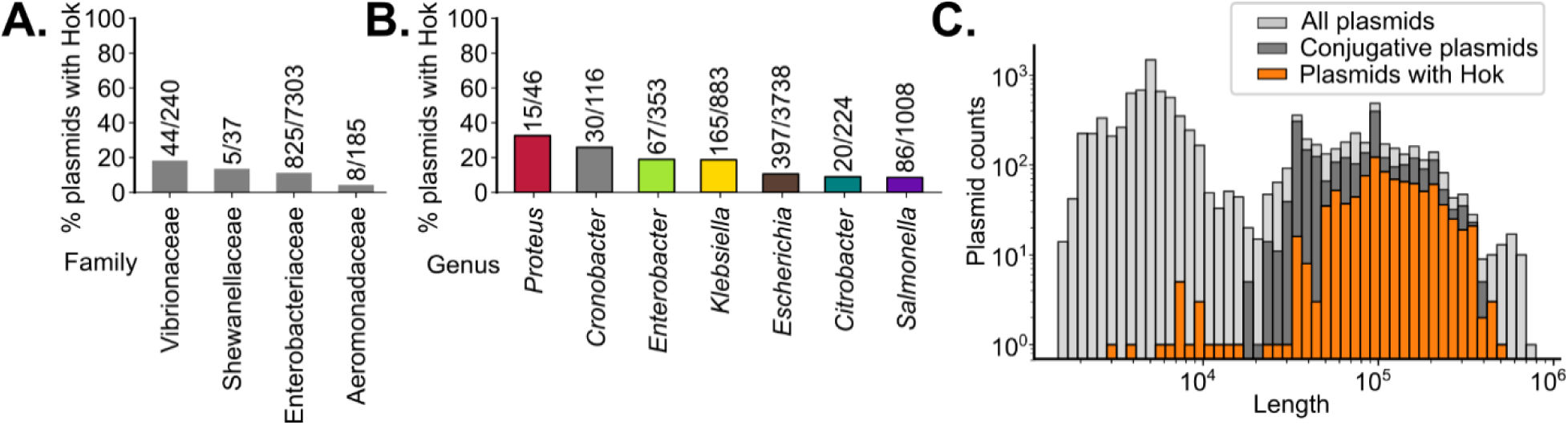
Hok loci are enriched in large plasmids. **A.** Percentage of plasmids containing *hok* per bacterial family**. B.** Percentage of plasmids with *hok* in Enterobacteriaceae genera where at least 10 *hok-*containing plasmids were identified. The number of plasmids containing a *hok* loci over the total number of plasmids are indicated on panels A and B. **C.** Histogram of the distribution of plasmid length for all plasmids from the IMG_PR database (light grey), conjugative plasmids (dark grey) or Hok-encoding plasmids (orange).

Then, we wondered whether the size of the plasmids was correlated with the presence of *hok*. For this we only considered plasmids form Enterobacteriaceae because most of the homologs were found in this family. The presence of *hok* was strongly associated with the plasmid size. While 65% of plasmids in Enterobacteriaceae are less than 25 kb, only 1.97% of the plasmids with Hok homologs were found in this size range (Fig. 6C). The size of *hok*-containing plasmids highly overlaps with the size of conjugative plasmids. Indeed, 99% of conjugative plasmids are longer than 25 kb (Fig. 6C). When considering only plasmids that contained at least one homolog, we did not observe any correlation between plasmid length and number of Hok homologs (Pearson coefficient=0.026, p-value=0.4). Thus, although *hok* loci are preferentially found in large plasmids, longer plasmids do not necessarily contain more homologs.

The presence of TAs has been correlated with the presence of partitioning systems in plasmids (47). To test if this is also the case for Hok, we looked for proteins involved in partitioning in the plasmid annotation. As expected, we mostly found partitioning proteins in plasmids longer than 25 kb (similar to the size of *hok*-containing plasmids (Fig. 6C). Partitioning proteins were present in 69% of plasmids longer than 25 kb, 37% of which also encoded Hok. In contrast, only 11% of the plasmids without partitioning systems encode a Hok (considering only plasmids longer than 25 kb). This suggests that *hok* is indeed enriched in plasmids with Par proteins.

### Inter-genera horizontal gene transfer of Hok is mainly mediated by plasmids

As observed in figure 1C, highly abundant 60% identity clusters tend to contain sequences found in more than one genus (Fig. 1C, Fig. S7), which suggests that Hok homologs could be transferred horizontally across genera. To further investigate this phenomenon, we looked at identical protein sequences present in multiple genera and identified 5.7% of the unique sequences in more than one genus. To investigate possible mechanisms mediating HGT, we compared the localization of all unique sequences with the localization of the sequences shared by more than one genus. Looking at the total number of unique peptides, most unique sequences (51%) were encoded only in chromosomes, 25% only in plasmids, 6.6% in prophages and 4.2% in phages (Fig. 7). The remaining 13% was found in more than one localization mostly shared between prophages, phages and chromosomes and only very rarely shared between chromosomes and plasmids (2.2%) (Fig. 7). In contrast, 72% of the peptides in the shared fraction were encoded in plasmids or both on plasmids and chromosomes, suggesting that most of the inter-genera HGT of Hok is mediated by plasmids (Fig. 7). To investigate the additional vehicles present on genomes that would enable HGT, we analyzed the genomic context of chromosomal loci containing Hok sequences that were either exclusively chromosomal or shared between chromosomes and plasmids. For that we first reduced sequence redundancy by clustering the loci based on average nucleotide identity over 20 kb adjacent to *hok.* Then, we visualized one representative locus from each cluster and colored the genes by annotated protein function (Fig. S9). Our analysis retrieved a high number of proteins related to insertion sequences (IS elements) and phages in loci with Hok homologs present also in plasmids (Fig. S9). We also observed some instances of plasmidic proteins (such as plasmid mobilization protein MobA and Plasmid SOS inhibition protein A), suggestive of plasmid integration events. In contrast, loci containing Hok sequences not found on plasmids predominantly contained phage-related proteins, with only infrequent occurrences of integrative and conjugative elements (ICEs) or IS elements (Fig. S9).

**Figure 7.**
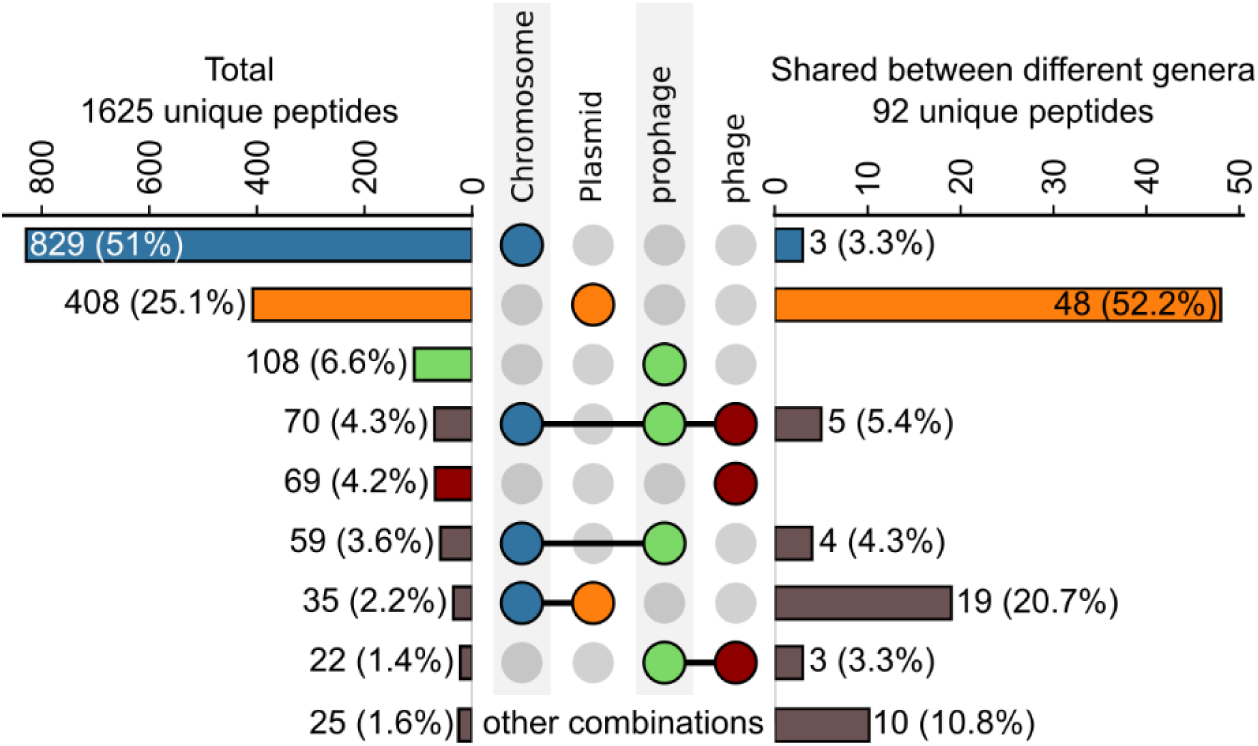
Inter-genera *hok* horizontal gene transfer is mostly mediated by plasmids. Comparison of total unique peptides (n=1625) vs. unique peptides shared across genera (n=92) found on chromosomes, plasmids, prophages and phages. Individual dots represent the presence of sequences only in one molecule type, while connected dots indicate sequences shared between those molecule types. Number and percentage from the total unique peptides are shown. Intersections representing less than 1% of the sequences were aggregated in the bars labelled “other combinations”.

Overall, Hok unique sequences were shared by up to 15 different genera (Fig. S8). The highest number of sequences were shared among *Escherichia, Salmonella*, *Klebsiella*, *Enterobacter* and *Citrobacter* (Fig. S8); in particular between *Escherichia* and *Salmonella* (43), and between *Escherichia* and *Klebsiella* (33) (Fig. S8). Together, these data show that detection of transferred (unmutated) sequences between plasmids and chromosomes is rare, either because such transfer does not occur or because acquired sequences rapidly mutate and become untraceable. In addition, inter-genus horizontal gene transfer is mostly mediated by plasmids. Sequences belonging to three 60% identity clusters (47, 15 and 35), which were mostly composed of plasmidic sequences, accounted for 74% of the shared sequences with cluster 47 containing almost half of all shared sequences (46.7%) (Fig. S8).

## MATERIAL & METHODS

### Search of Hok homologs in the RefSeq database

All complete bacterial genomes were downloaded from the NCBI RefSeq database (accessed on the 27^th^ October 2022), including 29655 assemblies (28). A list of 87 unique Hok sequences from the T1TAdb (release 6) served as an input for three independent similarity searches using MMseqs2, blastn and Infernal CMSearch (24). The hits were compiled in Table S1 with information about the protein sequence, nucleotide sequence and bacterial taxonomic classification.

MMseqs2 (version 11.e1a1c) was used to predict Hok homologs based on the peptide sequence (30). The peptides from the T1TAdb were first manually curated to remove very divergent or wrongly annotated sequences (TA06130, TA06119, TA06165 and TA05988). In addition, two sequences that were predicted by Fozo et al. 2010 but absent from the T1TAdb were included (RefSeq_start-stop coordinates: NC_007946.1_1566825_1567316 and NC_009425.1_87429_87587) (21). Then, the similarity search was performed using a minimum amino-acid length of 15 and 5 iterations.

The blastn search was performed using a set of unique predicted *hok* mRNA sequences from the T1TAdb using BLAST, version 2.12.0+ (29). The following parameters, similar to the ones reported in Fozo et al., 2010, were used: PAM70 matrix, wordsize of 2, no composition based statistics, a maximum number of targets of 1×10^6^, gapcost=10 and extension=2. Only hits with >80% identity (number of nucleotide matches divided by total number of nucleotides) were kept.

A search based on RNA sequence and secondary structure covariation was done using Infernal version 1.1.4 (31). Covariance models were built with the Infernal CMbuild module with default parameters. A first covariance model (CM1) was computed with the sequence and structure of full-length *hok* mRNA from the R1 plasmid (T1TAdb ID TA07123) (22). CM1 was used to scan the RefSeq database with the Infernal CMsearch tool. A second model (CM2) was constructed with the 101 highest non-redundant scoring hits that were then aligned using Infernal CMalign tool and used to scan once more the RefSeq database (Files S5 and S6). Before performing the search, the covariance models were calibrated using Infernal CMcalibrate module with the default parameters. To assess the contribution of RNA secondary structure to the search, we reconstructed the covariance model using the same alignment as for CM2, but with the --noss option in CMbuild, which ignores secondary structure annotations in the input alignment. Sequences with at least 300 nt were kept. The NCBI annotation for each of the identified loci (50 nt upstream and 200 nt downstream of the initial alignment) were downloaded and the CDS annotation (’NCBI_product’, ‘NCBI_protein_id’, ‘NCBI_translation’, ‘NCBI_gene’) was retrieved if the product contained any of the following words ‘Hok’,’Mok’,’Gef’ or ‘Sok’.

### Hok peptide prediction and filtering

To obtain complete putative Hok CDS, we extracted 50 nt upstream the start and 200 nt downstream the end of each hit. The Hok peptide sequences found with MMseqs2 were used to annotate the CDS from the blastn and CM searches. For that, we first identified sequences that were identical to any of the predicted peptides by searching the forward frames. If an identical match was not found, we aligned the translated sequences to each of the known peptides using Biopython 1.83 pairwise2.align.localms (gap-open penalty -2 and gap-extension penalty of -1) and retrieved the one with the highest score. Then the CDS was extended to the first start and stop codons upstream and downstream, respectively. Some of the sequences found by MMseqs2 could potentially include regions upstream the real start codon, causing the predicted CDS to extend to an upstream start codon, including regions that are not part of the natural peptide. To prevent this, we aligned the peptides using Muscle5 and kept the starting residue only if its corresponding column in the alignment had less than 90% of gaps. From this trimmed peptide, we predicted the closest upstream Shine-Dalgarno sequence (SD, GGAGG with at most 2 substitutions or indels) followed by maximum 10 nt and the start codon (ATG, GTG or TTG). Sequences with predicted SD and start codons were labeled ‘RBSstart’ in the ‘reannot_method’ column (Table S1). If we failed to identify a SD, we extended the amino-acid sequence upstream until a start codon (ATG, GTG, TTG) was found (marked as ‘start’ in the ‘reannot_method’ column). The CDS was extended downstream until a stop codon (TAA, TGA, TAG) was found. When the start codon could not be identified, the original hit alignment sequence was kept (added in the ‘pred_cds’ column and marked as ‘aln’ in the ‘reannot_method’ column). The position of the SD and start relative to the beginning of the extracted nt sequences were added to the table. When the same locus was identified by more than one method (*i.e.* contained overlapping CDS), only one entry was kept. The search methods that identified each homolog were recorded in the ‘Predicted_by’ column.

In order to remove false positives and degenerated sequences we excluded sequences that were smaller than 40 amino-acids or contained internal stop codons. In addition, we aligned the peptides using Muscle5, removed columns with more than 90% gaps using trimAl and kept only sequences longer than 40 residues aligning in conserved positions (using SeqKit) (48). In addition, we used CLANS to remove sequences that were not connected with the main cluster using a p-value of 1×10^-4^ as a threshold to connect two nodes (Fig. S1) (32).

### Peptide clustering and diversity analysis

To determine the diversity of Hok peptides, we computed an all-against-all identity matrix performing an MMseqs2 search with the non-redundant predicted peptides as both subject and query using easy-search with the following parameters: --prefilter-mode 2, --max-seqs 1000000, -s 7.5, --num-iterations 10, --mask 0, -c 0.8, --cov-mode 0. The resulting matrix was used to calculate the distribution of identities across all predicted homologs.

We also clustered the predicted CDS using MMseqs2 easy-cluster with high sensitivity (-s 7.5), minimum sequence identity of 60% (--min-seq-id 0.6) and 50 kmers per sequence (--kmer-per-seq 50). A sequence logo was generated to visualize amino-acid conservation by aligning non-redundant sequences with Muscle5 (version 5.3.linux64, -super5 algorithm), trimming with trimal -gt 0.10 to remove columns in the alignment with more than 90% gaps and plotting using LogoMaker version 0.8 (49, 50). Conservation of specific codons was observed by taking the codons corresponding to the alignment above and plotting the proportion of each nucleotide per position using LogoMaker. Only non-redundant CDS sequences were considered.

### Bacterial cultures

Bacterial strains are listed in Table S3. Cloning was done in *E. coli* Top10. Hok toxicity experiments were conducted in *Escherichia coli strain BW25133*. Bacteria were grown on LB supplemented with chloramphenicol (Cm) or Erythromycin (Erm) when needed at a concentration of 30 µg/mL and 300 µg/mL respectively. Liquid cultures were incubated at 37 °C with shaking at 200 rpm in flasks and 600 rpm in 96-well plates except when indicated otherwise.

### Construction of plasmids for toxicity assays

Plasmids used in this study are listed in Table S4. Experimentally tested Hok sequences are listed in Table S2. We selected several Hok representatives from the 16 most abundant clusters as well as from clusters with only one sequence. From the 26 clusters containing only 1 Hok sequence, we selected the 14 sequences with a length between 48 and 55 amino-acids and starting with a methionine. We mutated conserved residues to alanine. All the Hok CDS were amplified with Phusion polymerase (Thermo Scientific™) from synthetic DNA (gBlocks and eBlocks from IDT, sequences listed in (Table S2) except for Hok R1, Hok R1*Stop (adding a stop codon), HokB, HokA, and HokE that were amplified from genomic DNA of *E. coli* MG1655 or LR27 (Table S3). The specific primers used are listed in Table S5. First, the *hok*/Sok homolog loci TA05999 (24) was amplified with LRO3 and LRO228 and inserted in pAZ3 by digestion with EcoRI and HindIII (Thermo Scientific™) followed by ligation using T4 DNA ligase (Thermo Scientific™) to construct LRP38. All other plasmids were constructed by a modified AQUA cloning protocol (51) using LRP38 as a backbone. For that, LRP38 was cut with EcoRI and PCR-amplified using LRO491/608. PCR products were purified using the EasyPure® PCR Purification Kit and incubated with a seven-fold molar excess of insert in a final volume of 10 μL for 1 h at room temperature in a 96-well plate. Fifty microliters of *E. coli* TOP10 competent cells were transformed with the mixture via heat shock (1 minute at 42 °C followed by 2 minutes on ice). The bacteria were recovered for 1h at 37°C in LB containing 0.2% glucose. Transformants were plated in LB agar with Cm and glucose 0.2%. The sequence of the insert was checked by Sanger sequencing after colony PCR amplification with LRO273/LRO497 (Dream Taq polymerase, Thermo Scientific™). We did not obtain clones (or only mutated clones) for the following CDS (clus100_clus60): 15_1423 (PndA from R16), 39_851 and 39_866, 66_1506, 76_1723 (SrnB), that were therefore not tested for toxicity. The plasmids carrying CDS 19_179, 60_1482 and 69_1540, the only non-toxic Hok sequence, were sent for full plasmid sequencing and did not show any mutations compared to the control plasmid.

### Toxicity drop assays

Thermocompetent cells of *E. col*i BW25113 containing the pJAT13*araE* plasmid were transformed with each of the *hok* CDS overexpression plasmids. Single transformants were incubated in 150 µL of LB containing Cm, Erm and 0.2% glucose in 96-well plates and incubated overnight at 30°C, with 200 rpm agitation. Ten-fold serial dilution in 90 µl of LB until dilution 10^-7^ were done using the ASSIST PLUS pipetting robot (Integra Biosciences) and plated on Cm/Erm plates supplemented with 0.2% glucose or 0.01% arabinose and incubated at 37 °C during 24 h. The plates were imaged using the Amersham™ ImageQuant™ 800 from Cytiva. These experiments were repeated three times.

### Bacterial phylogenetic analysis

The bacterial phylogenetic tree was constructed using the Genome Taxonomy Database Toolkit (GTDB-Tk v2.4.0) using the ‘de_novo_wf’ workflow using p Chloroflexota as an outgroup and the --bacteria argument (52). The initial tree included all chromosomes in which the search was conducted in addition to the genomes in T1TAdb (even if they were not found in the RefSeq database). Plasmids were not included. The tree was then converted to iTOL-compatible format using gtdbtk convert_to_itol, uploaded to iTOL v6 and pruned to include only the most abundant genera of the Enterobacteriaceae family (for which we downloaded more than 100 chromosomes) (53). The classification assigned by GTDB-Tk was used for the distribution analyses.

### Co-localization of hok genes in genomic islets

We define a genomic islet as a region containing two or more *hok* homologs located within 2 kb of each other on the same chromosome or plasmid.

### Colocalization of Hok loci with defense systems

Phage defense systems were predicted in all genomes where Hok was found using DefenseFinder version 1.3.0 (41). Only replicons containing at least one defense system and one Hok homolog were considered for downstream analysis. To determine if *hok*/Sok systems were enriched in the vicinity of defense systems we calculated the percentage of Hok homologs that were less than 20 kb apart from a defense system (other than *hok*/Sok). This was compared with the probability of having Hok in the vicinity of a defense system if the localization was random. For this, we randomized the coordinates of the predicted homologs in each replicon (keeping the lengths and number of homologs unchanged) and calculated the proportion that fell <20 kb from any defense system. The randomization process was repeated 10 times to calculate the standard deviation. We repeated the process using SymE homologs, which belong to another T1TA family previously found to colocalized with restriction modification systems (42). For this, SymE proteins were predicted in the replicons were Hok was identified, as explained in the search section, but using only MMseqs2 and the annotated SymE peptides in the T1TAdb. Randomization was also performed as above.

### Identification of Hok homologs in prophages from RefSeq genomes

Sequences of phages were obtained from the IMG_VR virus database (version v4.1, directory 2022-12-19_7.1, containing phages predicted with high-confidence) (45). To identify Hok-containing prophages in RefSeq genomes, we mapped the prophages from the IMG_VR on the assemblies with Hok homologs. The unique phage identifier (UVIG) of the mapped phages was included in Table S1. Hok was considered encoded in the prophage if the positions of the phage and the Hok locus overlapped by at least 1 nucleotide. The percentage of Hok homologs encoded in prophages was calculated for the replicons containing at least one of each. Randomization was done as before to calculate the expected probability of a Hok homolog landing within a prophage if their distribution was random in the replicon. For comparison, the analysis was repeated for SymE homologs.

### Searches of Hok homologs in plasmids and phages

We also searched for Hok homologs in plasmids and phages in dedicated databases. Plasmids were obtained from the IMG_PR plasmid database (version 2023-08-08_1)(46). Then, we performed MMseqs2 (30) searches with default parameters (without iterations) in the nucleotide files of the IMG_PR and IMG_VR databases (IMGPR.nucl.fna and IMGVR_all_nucleotides-high_confidence.fna, respectively). As input, we used the predicted Hok peptides that resulted from the search in the RefSeq database after removing short and non-conserved sequences. For this, the unique Hok sequences were aligned with Muscle5 (version 5.3.linux64) using the -super5 argument and columns with low conservation (>90% gaps) were removed using trimal -gt 0.10 (54). After, sequences that were shorter than 40 amino-acids were removed using SeqKit2 seq -m 39 followed by duplicate removal using SeqKit (Version: 2.1.0) rmdup (55). Duplicated hits were removed if they overlapped by at least 1 nt, keeping the one with the lowest e-value given by MMseqs2. The CDS of the new sequences was predicted as described above. Sequences for which we could not find a start codon in the 50 nt upstream were also removed. The plasmids were marked as conjugative according to the annotation table in the PlasmidScope database (56). Similarly, we use the protein annotations in this database to identify plasmids with partitioning proteins if the product name contained one of the following terms: ParB, ParA, ParG, KorB, ParM, SopB, StbC, ParF, IncC, SopA, StbA, ParR, StbB and partition, retrieved from (47). Hits in the IMG_VR database were only kept if a prophage with the same UVIG was not previously identified in the RefSeq sequences. The new hits were added to Table S1. In Table S1 the RefSeq accession for hits retrieved from the IMG/PR database corresponds to the closest reference, defined as the reference plasmid showing the best alignment to the query sequence.

### Horizontal gene transfer analysis

Identical Hok sequences (100% identity clusters) that were found in more than one genus were selected and their genomic context was analyzed. We focused on loci located in the chromosome with Hok sequences either encoded in chromosomes only or encoded both in chromosomes and plasmids. We first downloaded the nucleotide sequences spanning from 10 kb upstream to 10 kb downstream of the *hok* homologs from the RefSeq database. We then reduced sequence redundancy by clustering based on the average nucleotide identity (ANI). For this, we constructed a similarity matrix by performing all-against-all pairwise ANI calculations using FastANI (version 1.33 with a fragment length of 500 nt) (57). Sequences with least 80% ANI over at least 10 mapped fragments (corresponding to 5 kb), were considered to be in the same cluster. Sequences with the smallest average distance to all other elements in the cluster (the highest average ANI) were selected as cluster representatives. The cluster representatives were then visualized by clinker (58). To standardize and categorize gene annotations, we applied a series of string-matching filters to the product annotation provided by clinker. Each entry was evaluated for the presence of specific keywords or patterns indicative of mobile genetic elements. Matching entries were reassigned to curated category labels for consistency.

The criteria for the classification were as follows:

1. Plasmid Identification: Entries containing the term *“plasmid”* were assigned the label “plasmid”.
2. Insertion Sequences (IS): Products starting with the prefix *“IS”* were labeled as “IS”.
3. Phage-Related Genes: Entries containing phage-associated terms including *phage, tail, holin, Cro/CI, lysis, lysozyme, superinfection, terminase, excisionase* or *capsid* were grouped under the label “phage-related”.
4. Integrative and Conjugative Elements (ICE): Entries containing *“ICE”* or *“conjugative element”* were categorized as “ICE”.
5. Conjugation Genes: Entries containing the substring *“conj”* were labeled as “conjugation”.
6. Transposases: Entries containing *“transposase”* or beginning with *“Tn”* were assigned the label “transposase”.
7. Integrases and Recombinases: Entries with *“integrase”* or *“recombinase”* were labeled accordingly.
8. Toxin-Antitoxin Systems: Genes containing *“toxin-antitoxin”*, *“GnsA/GnsB”*, *“YdaT”*, or *“YdaS”* were labeled as “TA”.

All filtering steps used regular expression matching (regex=True) and were case-insensitive.

### Data analysis and visualization

Results were analyzed and plotted using Jupyter Notebook (Python 3.8, Pandas 2.0.3, matplotlib3.7.3, seaborn 0.13.2 and upsetplot 0.9.0) unless indicated otherwise.

## DISCUSSION

Prediction of T1TA homologs has been notoriously difficult due to the small size, low conservation and lack of catalytic domains of most of the toxins (7, 21). To ensure retrieving the highest number of homologs, we applied a combination of strategies using protein sequence, RNA sequence, or RNA sequence and structure, to search against the complete bacterial genomes of the RefSeq database. We identified 32,532 Hok homologs in 59 bacterial genera. Most of the loci we retrieved were already annotated as Hok in this database, reflecting the fact that Hok had been thoroughly studied and validating our search methods. Limiting our search to complete genomes allowed us to calculate the total number of homologs per chromosome and assess their co-localization with phage defense systems and prophages. However, this also reduced the diversity of the analyzed sequences by excluding difficult-to-assemble metagenomes and biasing the sample towards more studied culturable bacteria. Of note, the searches in the plasmid and phage databases included partially completed sequences and metagenomic data.

RNA structure is important for regulating T1TA expression and has been previously used for finding *hok* homologs (7, 22). Of the three search tools used in this work, Infernal (which incorporates RNA structure) returned the highest number of hits. Even though the majority of the Hok homologs could also be retrieved by Infernal without taking RNA structure into account, 16% of the unique peptides were only found when the predicted structure was used. In addition, there were 9 identity clusters that were detected by Infernal only when the RNA structure was used, which strengthens the case for using RNA structure for increasing the diversity of predicted T1TA homologs.

The *hok*/Sok system contains multiple regulatory elements that are essential for its function. These include the Sok small RNA antitoxin, the Mok leader peptide and the structure of the *hok* mRNA. Due to the difficulties associated with the correct identification of these elements, we focused on the analysis of the Hok coding sequence. Indeed, trying to identify the Mok coding sequence is not trivial as predicting putative start codons, along with their associated SD in the three reading frames, leads to several possibilities. Similarly, *hok* mRNA and Sok sequences were previously bioinformatically predicted for the ones in the T1TAdb but experimental validation would be needed to confirm the exact 5 and 3’ ends (24). Future studies could take advantage of the identified peptide homologs to further investigate the characteristics of each of these elements.

In 2017, Coray et al. reported that the distribution of T1TAs was limited compared to other TAs (1). However, more genome sequences have become available, along with new databases and advanced search tools. Consistent with previous results, we found that Hok is restricted to 4 families in the Pseudomonadota phylum (previously Proteobacteria) (1). Thus, the addition of new sequences and the combination of different search strategies did not expand the distribution of Hok. It is however intriguing that the prevalence of Hok is so low in the other families compared to Enterobacteriaceae. Furthermore, while the percentage of chromosomes with *hok* is much higher than the percentage of *hok*-containing plasmids in Enterobacteriaceae, the opposite is true for the rest of the families (Fig. 3A). It remains to be determined whether these patterns arise from the high genomic redundancy in Enterobacteriaceae, or whether it reflects ecological or physiological differences between the families, such as the differences in the exposure to phages or in Hok toxicity in these bacteria. It is also possible that the observed low prevalence of Hok in these families is an artifact due of the difficulty to detect more distant homologs.

We highlighted the high sequence variability within Hok peptides by obtaining a total of 79 clusters with 60% identity (61 clusters from the RefSeq database and 18 clusters from the plasmid and phage database) (Fig. S7). This variability was already featured in the T1TAdb which spanned 23 of the 60% identity clusters (Fig. 1C and S7). Our bioinformatic searches, with 51 novel 60% clusters, contributed to expand this diversity. Because the high sequence diversity prevented the construction of a reliable phylogenetic tree, clustering was used as an alternative strategy to identify and interpret patterns of sequence variability. However, even if some evolutionary relationships might be preserved in the clusters, there is currently no reliable method to evaluate their monophyly or infer the directionality of sequence divergence. As such, we could not draw any conclusions about the origin and evolutionary relationships of these clusters.

At the amino-acid sequence level, we observed that only 7 out of the 54 aligned positions (13%) were conserved in more than 80% of the sequences (Fig. 1). However, we show here that these amino-acids are individually not strictly required for toxicity as individually mutating those residues to alanine does not abrogate toxicity, at least when Hok is overexpressed (Fig. 2). Apart from the role of the two conserved cysteines, nothing is known about the importance of other residues in the activity of Hok (8, 9, 25). The limited biochemical knowledge available on Hok makes it difficult to predict activity and understand the origin and function of its sequence diversity. For example, while most of the sequence diversity was found in chromosomes, it is possible that many of these sequences are inactive, in the process of degeneration or have evolved other functions. Indeed, previous reports have suggested that chromosomal Hok homologs are mostly inactive, at least in the laboratory *E. coli K12* strain (26). However, another hypothesis is that chromosomal TA could defend the host against plasmid addiction caused by plasmidic TAs (59, 60). Here, we show that Hok peptides originating from chromosomes are toxic under overexpression. Our analysis also showed similar toxicity profiles for Hok CDS from both high and low abundant clusters (Fig. S4). Note that our experimental design, which involves CDS overexpression from a multicopy plasmid, may not accurately reflect toxicity differences that could be relevant under natural conditions. Indeed, in addition to native copy numbers and promoter strengths, the full mRNA would need to be expressed in order to include other regulatory elements influencing the toxicity of the different homologs. Therefore, sequences that showed toxicity under our experimental conditions are potentially functional, however we cannot exclude that there are differences in toxicity that are relevant under natural conditions. Whether the function of Hok in the chromosome is the same as in plasmids remains to be investigated. On the one side, the fact that we rarely found the same sequence in both argues towards a difference in function; a similar observation was done for plasmidic and chromosomal *cdd* systems leading to a proposed action in as anti-addictive modules (59). On the other, we failed to identify distinctive characteristics at the amino-acid level that would set them apart. Future mechanistic studies, together with comparative analyses like the one performed here, are expected to yield deeper insights into the evolution of Hok and the possible emergence of novel functions.

In light of the reported role of Hok in phage defense, we investigated the link between Hok, phages and prophages (15). We showed that *hok* was enriched in prophages. By mapping only highly-confidently predicted phages on chromosomes, we ensured we could accurately determine the prophage boundaries. However, this led to an underestimation of number of phages. Indeed, we only observed approximately 16% of chromosomes with prophages, while the literature reports up to 75% of bacteria containing prophages (61). The underestimation of the number of prophages likely leads to a systematic underestimation of the enrichment of *hok* within prophages. While the function of *hok* in prophages remains to be studied, multiple phages have been shown to encode defense systems, which could be a hint in this direction (44). To further study if Hok could be functioning as a phage defense system, we investigated the proximity of *hok* with other defense systems but did not observe any enrichment. It would also be interesting to analyze the correlation between number of *hok* genes and the number of prophages per chromosome. However, a confident prediction of all prophages in enterobacteria would be required to perform this analysis, which is outside the scope of this work.

Hok is a well-known actor of plasmid maintenance, and is used in the research and industrial setup for this purpose (62, 63). Here we describe that Hok is mainly found in large plasmids (> 25 kb) and its presence correlates with the presence of partitioning systems. Large plasmids have lower copy numbers per cell than small plasmids, are less likely to segregate passively and rely on active segregation mechanisms such as partitioning systems (64–66). Partitioning systems act together with TAs to avoid plasmid extinction in the population, explaining the presence of Hok to actively maintain those plasmids (47, 60). It was recently described that, comparably to anti-defense systems and other toxin-antitoxin systems, *hok*/Sok sequences are found in the leading region of conjugative plasmids, highlighting their possible role in early plasmid establishment (67).

While there is evidence that type II TAs are transferred horizontally, previous bioinformatic studies failed to find evidence for recent HGT between distant bacteria for two T1TA families (Ldr and Fst) (21, 68). Due to the short length of Hok, it is difficult to discern whether sequence similarities reflect recent divergence from a common ancestor or result from convergent evolution. Even a few mutations may cause vertically inherited sequences to appear highly divergent, potentially mimicking patterns expected from ancient divergence or horizontal acquisition from distant lineages. To minimize the impact of this confounding factors, we classified sequences as potentially horizontally acquired only if they were identical and detected across different genera. The peptides that were shared across genera represented only 5.7 % of the total unique sequences, however due to the stringency of our analysis this is likely an underestimation as it excluded sequences that diverged shortly after HGT. Nonetheless, these results provide a clear indication that long-distance HGT of Hok homologs is mainly mediated by plasmids. Indeed, while plasmids contain 25% of unique sequences, 53% of the shared unique sequences are found only in plasmids (Fig. 7). This is in agreement with observation that some plasmids are indeed transferred across genera (69). In addition, the genera in Enterobacteriacea that were reported to exchange more plasmids (*Escherichia, Salmonella, Klebsiella, Enterobacter* and *Citrobacter*), were also the genera that shared more identical sequences (69) (Fig. S8). This suggests that most of these reflect HGT events and not convergent evolution. In the few instances where identical sequences were found on chromosomes from different genera, they were carried by mobile genetic elements such as prophages and ICE (Fig. S9). Notably, most phages have a narrow host range (61, 70) and only a few exceptions of phages with the ability to infect more than one genus have been reported (71, 72). It would be valuable to investigate whether phages containing shared Hok sequences belong to broad range phages. Furthermore, the role of the different mobile genetic elements in *hok* HGT between closely related organisms remains to be explored. However, detecting HGT events within the same genus is more challenging and would require a more specialized approach.

In conclusion we present here the most comprehensive comparative genomics analysis on Hok to date, providing an exhaustive list of bacterial Hok sequences and localization. We assessed the toxicity of a wide range of homologs and provide a database of diverse active Hok sequences from the most abundant 60% clusters, arising both from plasmids and chromosomes, that could be further utilized for various applications such as plasmid stabilization or antimicrobial approaches (19, 20, 73). The information provided here may also pave the way for future studies aimed at understanding the function of Hok across different bacterial species and genomic localizations.

## DATA AVAILABILITY

The data underlying this article are available in the article and in its online supplementary material. The code used for data analysis and figure generation has been deposited on ttps://gitub.u-bordeaux.fr/alerhun/Escalera-Maurer_2025.

## ACKNOWLEDGEMENTS

We thank Fabien Darfeuille for plasmid gifts. pJAT13araE was a gift from Jay Keasling (Addgene plasmid # 18987; http://n2t.net/addgene:18987; RRID:Addgene 18987). We acknowledge Kira Makarova and Fabien Darfeuille for critical reading of the manuscript. We thank Marine Baraquin for technical help. We acknowledge members of the ARNA laboratory for helpful discussions.

## Author contributions

Andrés Escalera Maurer (Conceptualization, Data curation, Formal analysis, Investigation, Supervision, Visualization, Writing—original draft, Writing - review & editing), Adriana Messineo (Investigation, Visualization, Funding acquisition, Writing—review & editing), Thibaud T. Renault (Investigation, Supervision, Writing—review & editing), Elena Nicollin (Investigation), Erika Castaneda-Sastre (Investigation), Matthieu Brunot (Investigation), Cléo Berrehail (Methodology) and Anaïs Le Rhun (Conceptualization, Funding acquisition, Project administration, Supervision, Visualization, Writing—original draft, Writing—review & editing)

## References

1. Coray, D.S., Wheeler, N.E., Heinemann, J.A. and Gardner, P.P. (2017) Why so narrow: Distribution of anti-sense regulated, type I toxin-antitoxin systems compared with type II and type III systems. RNA Biol., 14, 275–280.

2. Harms, A., Brodersen, D.E., Mitarai, N. and Gerdes, K. (2018) Toxins, Targets, and Triggers: An Overview of Toxin-Antitoxin Biology. Mol. Cell, 70, 768–784.

3. Shore, S.F.H., Leinberger, F.H., Fozo, E.M. and Berghoff, B.A. (2024) Type I toxin-antitoxin systems in bacteria: from regulation to biological functions. EcoSal Plus, 0, eesp-0025-2022.

4. Bonabal, S. and Darfeuille, F. (2023) Preventing toxicity in toxin-antitoxin systems: An overview of regulatory mechanisms. Biochimie, 10.1016/j.biochi.2023.07.013.

5. Gerdes, K. and Wagner, E.G.H. (2007) RNA antitoxins. Curr. Opin. Microbiol., 10, 117–124.

6. Gerdes, K., Thisted, T. and Martinussen, J. (1990) Mechanism of post-segregational killing by the hok/sok system of plasmid R1: sok antisense RNA regulates formation of a hok mRNA species correlated with killing of plasmid-free cells. Mol. Microbiol., 4, 1807–1818.

7. Masachis, S. and Darfeuille, F. (2018) Type I Toxin-Antitoxin Systems: Regulating Toxin Expression via Shine-Dalgarno Sequence Sequestration and Small RNA Binding. Microbiol. Spectr., 6.

8. Bar-Yaacov, D., Mordret, E., Towers, R., Biniashvili, T., Soyris, C., Schwartz, S., Dahan, O. and Pilpel, Y. (2017) RNA editing in bacteria recodes multiple proteins and regulates an evolutionarily conserved toxin-antitoxin system. Genome Res., 27, 1696–1703.

9. Didi, L., Fargeon, O., Aspit, L., Elias, E., Braverman, D., Melamed, D., Keidar-Friedman, D., Sorek, N., Raz, O., Tamir, S.O., et al. (2025) A-to-I mRNA editing in bacteria can affect protein sequence, disulfide bond formation, and function. Nucleic Acids Res., 53, gkaf584.

10. Wilmaerts, D., Windels, E.M., Verstraeten, N. and Michiels, J. (2019) General Mechanisms Leading to Persister Formation and Awakening. Trends Genet. TIG, 35, 401–411.

11. Gerdes, K., Larsen, J.E. and Molin, S. (1985) Stable inheritance of plasmid R1 requires two different loci. J. Bacteriol., 161, 292–298.

12. Gerdes, K., Rasmussen, P.B. and Molin, S. (1986) Unique type of plasmid maintenance function: postsegregational killing of plasmid-free cells. Proc. Natl. Acad. Sci. U. S. A., 83, 3116–3120.

13. Gerdes, K., Helin, K., Christensen, O.W. and Løbner-Olesen, A. (1988) Translational control and differential RNA decay are key elements regulating postsegregational expression of the killer protein encoded by the parB locus of plasmid R1. J. Mol. Biol., 203, 119–129.

14. Fraikin, N. and Van Melderen, L. (2024) Single-cell evidence for plasmid addiction mediated by toxin-antitoxin systems. Nucleic Acids Res., 10.1093/nar/gkae018.

15. Pecota, D.C. and Wood, T.K. (1996) Exclusion of T4 phage by the hok/sok killer locus from plasmid R1. J. Bacteriol., 178, 2044–2050.

16. Verstraeten, N., Knapen, W.J., Kint, C.I., Liebens, V., Van den Bergh, B., Dewachter, L., Michiels, J.E., Fu, Q., David, C.C., Fierro, A.C., et al. (2015) Obg and Membrane Depolarization Are Part of a Microbial Bet-Hedging Strategy that Leads to Antibiotic Tolerance. Mol. Cell, 59, 9–21.

17. Cooper, T.F. and Heinemann, J.A. (2000) Postsegregational killing does not increase plasmid stability but acts to mediate the exclusion of competing plasmids. Proc. Natl. Acad. Sci., 97, 12643–12648.

18. Cooper, T.F., Paixão, T. and Heinemann, J.A. (2010) Within-host competition selects for plasmid-encoded toxin–antitoxin systems. Proc. R. Soc. B Biol. Sci., 277, 3149–3155.

19. Faridani, O.R., Nikravesh, A., Pandey, D.P., Gerdes, K. and Good, L. (2006) Competitive inhibition of natural antisense Sok-RNA interactions activates Hok-mediated cell killing in Escherichia coli. Nucleic Acids Res., 34, 5915–5922.

20. Chen, A., Dong, Y., Jiang, H., Wei, M., Ren, Y. and Zhang, J. (2024) Application of plasmid stabilization systems for heterologous protein expression in Escherichia coli. Mol. Biol. Rep., 51, 939.

21. Fozo, E.M., Makarova, K.S., Shabalina, S.A., Yutin, N., Koonin, E.V. and Storz, G. (2010) Abundance of type I toxin-antitoxin systems in bacteria: searches for new candidates and discovery of novel families. Nucleic Acids Res., 38, 3743–3759.

22. Steif, A. and Meyer, I.M. (2012) The hok mRNA family. RNA Biol., 9, 1399–1404.

23. Peng, J., Triplett, L.R., Schachterle, J.K. and Sundin, G.W. (2019) Chromosomally Encoded hok-sok Toxin-Antitoxin System in the Fire Blight Pathogen Erwinia amylovora: Identification and Functional Characterization. Appl. Environ. Microbiol., 85, e00724–19.

24. Tourasse, N.J. and Darfeuille, F. (2021) T1TAdb: the database of Type I Toxin-Antitoxin systems. RNA N. Y. N, 10.1261/rna.078802.121.

25. Wilmaerts, D., De Loose, P.-J., Vercauteren, S., De Smedt, S., Verstraeten, N. and Michiels, J. (2021) Functional analysis of cysteine residues of the Hok/Gef type I toxins in Escherichia coli. FEMS Microbiol. Lett., 368, fnab069.

26. Pedersen, K. and Gerdes, K. (1999) Multiple hok genes on the chromosome of Escherichia coli. Mol. Microbiol., 32, 1090–1102.

27. Nielsen, A.K., Thorsted, P., Thisted, T., Wagner, E.G. and Gerdes, K. (1991) The rifampicin-inducible genes srnB from F and pnd from R483 are regulated by antisense RNAs and mediate plasmid maintenance by killing of plasmid-free segregants. Mol. Microbiol., 5, 1961–1973.

28. O’Leary, N.A., Wright, M.W., Brister, J.R., Ciufo, S., Haddad, D., McVeigh, R., Rajput, B., Robbertse, B., Smith-White, B., Ako-Adjei, D., et al. (2016) Reference sequence (RefSeq) database at NCBI: current status, taxonomic expansion, and functional annotation. Nucleic Acids Res., 44, D733–745.

29. Camacho, C., Coulouris, G., Avagyan, V., Ma, N., Papadopoulos, J., Bealer, K. and Madden, T.L. (2009) BLAST+: architecture and applications. BMC Bioinformatics, 10, 421.

30. Steinegger, M. and Söding, J. (2017) MMseqs2 enables sensitive protein sequence searching for the analysis of massive data sets. Nat. Biotechnol., 35, 1026–1028.

31. Nawrocki, E.P. and Eddy, S.R. (2013) Infernal 1.1: 100-fold faster RNA homology searches. Bioinforma. Oxf. Engl., 29, 2933–2935.

32. Frickey, T. and Lupas, A. (2004) CLANS: a Java application for visualizing protein families based on pairwise similarity. Bioinforma. Oxf. Engl., 20, 3702–3704.

33. Gotfredsen, M. and Gerdes, K. (1998) The Escherichia coli relBE genes belong to a new toxin–antitoxin gene family. Mol. Microbiol., 29, 1065–1076.

34. Sakikawa, T., Akimoto, S. and Ohnishi, Y. (1985) Cloning and expression of the pnd gene of R16: determination of transcriptional direction and evolutionary analysis. Microbiol. Immunol., 29, 791–801.

35. Sakikawa, T., Akimoto, S. and Ohnishi, Y. (1989) The pnd gene in E. coli plasmid R16: nucleotide sequence and gene expression leading to cell Mg2+ release and stable RNA degradation. Biochim. Biophys. Acta, 1007, 158–166.

36. Loh, S.M., Cram, D.S. and Skurray, R.A. (1988) Nucleotide sequence and transcriptional analysis of a third function (Flm) involved in F-plasmid maintenance. Gene, 66, 259–268.

37. Makarova, K.S., Wolf, Y.I., Snir, S. and Koonin, E.V. (2011) Defense islands in bacterial and archaeal genomes and prediction of novel defense systems. J. Bacteriol., 193, 6039–6056.

38. Rocha, E.P.C. and Bikard, D. (2022) Microbial defenses against mobile genetic elements and viruses: Who defends whom from what? PLoS Biol., 20, e3001514.

39. Doron, S., Melamed, S., Ofir, G., Leavitt, A., Lopatina, A., Keren, M., Amitai, G. and Sorek, R. (2018) Systematic discovery of antiphage defense systems in the microbial pangenome. Science, 359, eaar4120.

40. Tesson, F., Hervé, A., Mordret, E., Touchon, M., d’Humières, C., Cury, J. and Bernheim, A. (2022) Systematic and quantitative view of the antiviral arsenal of prokaryotes. Nat. Commun., 13, 2561.

41. Tesson, F., Planel, R., Egorov, A.A., Georjon, H., Vaysset, H., Brancotte, B., Néron, B., Mordret, E., Atkinson, G.C., Bernheim, A., et al. (2024) A Comprehensive Resource for Exploring Antiphage Defense: DefenseFinder Webservice, Wiki and Databases. Peer Community J., 4.

42. Thompson, M.K., Nocedal, I., Culviner, P.H., Zhang, T., Gozzi, K.R. and Laub, M.T. (2022) Escherichia coli SymE is a DNA-binding protein that can condense the nucleoid. Mol. Microbiol., 117, 851–870.

43. Peltier, J., Hamiot, A., Garneau, J.R., Boudry, P., Maikova, A., Hajnsdorf, E., Fortier, L.-C., Dupuy, B. and Soutourina, O. (2020) Type I toxin-antitoxin systems contribute to the maintenance of mobile genetic elements in Clostridioides difficile. *Commun*. Biol., 3, 718.

44. Patel, P.H. and Maxwell, K.L. (2023) Prophages provide a rich source of antiphage defense systems. Curr. Opin. Microbiol., 73, 102321.

45. Camargo, A.P., Nayfach, S., Chen, I.-M.A., Palaniappan, K., Ratner, A., Chu, K., Ritter, S.J., Reddy, T.B.K., Mukherjee, S., Schulz, F., et al. (2023) IMG/VR v4: an expanded database of uncultivated virus genomes within a framework of extensive functional, taxonomic, and ecological metadata. Nucleic Acids Res., 51, D733–D743.

46. Camargo, A.P., Call, L., Roux, S., Nayfach, S., Huntemann, M., Palaniappan, K., Ratner, A., Chu, K., Mukherjeep, S., Reddy, T.B.K., et al. (2024) IMG/PR: a database of plasmids from genomes and metagenomes with rich annotations and metadata. Nucleic Acids Res., 52, D164–D173.

47. Effe, J., Santer, M., Wang, Y., Feenstra, T.E., Hülter, N.F. and Dagan, T. (2025) The combination of active partitioning and toxin-antitoxin systems is most advantageous for low-copy plasmid fitness. Nat. Commun., 16, 7078.

48. Capella-Gutiérrez, S., Silla-Martínez, J.M. and Gabaldón, T. (2009) trimAl: a tool for automated alignment trimming in large-scale phylogenetic analyses. Bioinformatics, 25, 1972–1973.

49. Edgar, R.C. (2022) High-accuracy alignment ensembles enable unbiased assessments of sequence homology and phylogeny. 10.1101/2021.06.20.449169.

50. Tareen, A. and Kinney, J.B. (2020) Logomaker: beautiful sequence logos in Python. Bioinforma. Oxf. Engl., 36, 2272–2274.

51. Beyer, H., Gonschorek, P., Samodelov, S., Meier, M., Weber, W. and Zurbriggen, M. (2015) AQUA Cloning: A Versatile and Simple Enzyme-Free Cloning Approach. PloS One, 10, e0137652.

52. Chaumeil, P.-A., Mussig, A.J., Hugenholtz, P. and Parks, D.H. (2019) GTDB-Tk: a toolkit to classify genomes with the Genome Taxonomy Database. Bioinforma. Oxf. Engl., 36, 1925–1927.

53. Letunic, I. and Bork, P. (2024) Interactive Tree of Life (iTOL) v6: recent updates to the phylogenetic tree display and annotation tool. Nucleic Acids Res., 52, W78–W82.

54. Edgar, R.C. (2022) Muscle5: High-accuracy alignment ensembles enable unbiased assessments of sequence homology and phylogeny. Nat. Commun., 13, 6968.

55. Shen, W., Le, S., Li, Y. and Hu, F. (2016) SeqKit: A Cross-Platform and Ultrafast Toolkit for FASTA/Q File Manipulation. PLOS ONE, 11, e0163962.

56. Li, Y., Feng, X., Chen, X., Yang, S., Zhao, Z., Chen, Y. and Li, S.C. (2025) PlasmidScope: a comprehensive plasmid database with rich annotations and online analytical tools. Nucleic Acids Res., 53, D179–D188.

57. Jain, C., Rodriguez-R, L.M., Phillippy, A.M., Konstantinidis, K.T. and Aluru, S. (2018) High throughput ANI analysis of 90K prokaryotic genomes reveals clear species boundaries. Nat. Commun., 9, 5114.

58. Gilchrist, C.L.M. and Chooi, Y.-H. (2021) clinker & clustermap.js: automatic generation of gene cluster comparison figures. Bioinforma. Oxf. Engl., 37, 2473–2475.

59. Saavedra De Bast, M., Mine, N. and Van Melderen, L. (2008) Chromosomal toxin-antitoxin systems may act as antiaddiction modules. J. Bacteriol., 190, 4603–4609.

60. Brendler, T., Reaves, L. and Austin, S. (2004) Interplay between Plasmid Partition and Postsegregational Killing Systems. J. Bacteriol., 186, 2504–2507.

61. López-Leal, G., Camelo-Valera, L.C., Hurtado-Ramírez, J.M., Verleyen, J., Castillo-Ramírez, S. and Reyes-Muñoz, A. (2022) Mining of Thousands of Prokaryotic Genomes Reveals High Abundance of Prophages with a Strictly Narrow Host Range. mSystems, 7, e0032622.

62. Srivastava, A., Pati, S., Kaushik, H., Singh, S. and Garg, L.C. (2021) Toxin-antitoxin systems and their medical applications: current status and future perspective. Appl. Microbiol. Biotechnol., 105, 1803–1821.

63. Lin, J., Guo, Y., Yao, J., Tang, K. and Wang, X. (2023) Applications of toxin-antitoxin systems in synthetic biology. Eng. Microbiol., 3, 100069.

64. Ramiro-Martínez, P., Quinto, I. de, Lanza, V.F., Gama, J.A. and Rodríguez-Beltrán, J. (2025) Universal rules govern plasmid copy number. 10.1101/2024.10.04.616648.

65. Rodríguez-Beltrán, J., DelaFuente, J., León-Sampedro, R., MacLean, R.C. and San Millán, Á. (2021) Beyond horizontal gene transfer: the role of plasmids in bacterial evolution. Nat. Rev. Microbiol., 19, 347–359.

66. Sengupta, M. and Austin, S. (2011) Prevalence and Significance of Plasmid Maintenance Functions in the Virulence Plasmids of Pathogenic Bacteria ▿. Infect. Immun., 79, 2502–2509.

67. Samuel, B., Mittelman, K., Croitoru, S.Y., Ben Haim, M. and Burstein, D. (2024) Diverse anti-defence systems are encoded in the leading region of plasmids. Nature, 635, 186–192.

68. Ramisetty, B.C.M. and Santhosh, R.S. (2016) Horizontal gene transfer of chromosomal Type II toxin–antitoxin systems of Escherichia coli. FEMS Microbiol. Lett., 363, fnv238.

69. Redondo-Salvo, S., Fernández-López, R., Ruiz, R., Vielva, L., de Toro, M., Rocha, E.P.C., Garcillán-Barcia, M.P. and de la Cruz, F. (2020) Pathways for horizontal gene transfer in bacteria revealed by a global map of their plasmids. Nat. Commun., 11, 3602.

70. Piel, D., Bruto, M., Labreuche, Y., Blanquart, F., Goudenège, D., Barcia-Cruz, R., Chenivesse, S., Le Panse, S., James, A., Dubert, J., et al. (2022) Phage-host coevolution in natural populations. Nat. Microbiol., 7, 1075–1086.

71. Schwarzer, D., Buettner, F.F.R., Browning, C., Nazarov, S., Rabsch, W., Bethe, A., Oberbeck, A., Bowman, V.D., Stummeyer, K., Mühlenhoff, M., et al. (2012) A multivalent adsorption apparatus explains the broad host range of phage phi92: a comprehensive genomic and structural analysis. J. Virol., 86, 10384–10398.

72. Matsuzaki, S., Tanaka, S., Koga, T. and Kawata, T. (1992) A broad-host-range vibriophage, KVP40, isolated from sea water. Microbiol. Immunol., 36, 93–97.

73. Danino, T., Prindle, A., Kwong, G.A., Skalak, M., Li, H., Allen, K., Hasty, J. and Bhatia, S.N. (2015) Programmable probiotics for detection of cancer in urine. Sci. Transl. Med., 7, 289ra84–289ra84.

