## supplementary data for "The Hok bacterial toxin: diversity, toxicity, distribution and genomic localization"

Andrés Escalera-Maurer1 (ORCID 0000-0002-2660-262X), Adriana Messineo1 (ORCID 0009-0004-8557-9863), Thibaud T. Renault1 (ORCID 0000-0002-1530-2613), Elena Nicollin1 (ORCID 0009-0002-4882-9590), Erika Castaneda-Sastre1 (ORCID 0000-0002-4411-753X), Matthieu Brunot1, Cléo Berrehail1 (ORCID 0009-0005-6920-3839) and Anaïs Le Rhun1* (ORCID 0000-0003-2211-2153)

---

**This PDF file includes:**

**Legends of Files S1 to S6 and of Table S1 and S2 (Separate files)**

**Legends and Fig. S1 to S9**

**Legends and Tables S3 to S5**

**Supplementary references**

---

**Supplementary File S1 (separate file)**
Protein alignments using Muscle5 of homologs found only in chromosomes displaying conserved positions with less than 90% gaps across aligned sequences. Used in Fig. S3B.

**Supplementary File S2 (separate file)**
Protein alignments using Muscle5 of homologs found only in plasmids displaying conserved positions with less than 90% gaps across aligned sequences. Used in Fig. S3B.

**Supplementary File S3 (separate file)**
Alignments of all unique protein sequences using Muscle5 displaying conserved positions with less than 90% gaps across aligned sequences. Used in Fig. 1.

**Supplementary File S4 (separate file)**
Protein alignments using Muscle5 of homologs in the Vibrionaceae family displaying conserved positions with less than 90% gaps across aligned sequences. Used in Fig. S6B.

**Supplementary File S5 (separate file)**
This file contains the covariance model (.cm) for the *hok* mRNA family built with the top 101 sequences obtained by our first search.

**Supplementary File S6 (separate file)**
This file contains the multiple sequence alignment of 101 *hok* mRNA homologs in Stockholm format (`.sto`), used to build the covariance model.

**Table S1 (separate file)**

**Summary of all hits obtained by the similarity searches in the refseq, IMG_PR and IMG_VR database.**

The columns are: **refseq_acc**, replicon accession number. For the hits retrieved from the IMG_PR db refseq the accession corresponds to the closest reference (The reference plasmid that best aligned with the sequence: the minimum average nucleotide identity between the plasmid and its closest reference. was 79% with at least 50% of the plasmid's length aligned to the reference sequence) ; **start** and **end,** coordinates of the initial hit; **clus100**, identifier for each unique predicted peptide sequence: **clus60**, 60% identity cluster to which each predicted peptide belongs; **removed**, hits removed and removal method; **prediction_by**, identification method for each hit; **UVIG**, identifier of the Uncultivated Viral Genome in the IMG_VR database; **plasmid_id**, identifiers as per the IMG_PR database; **locus_id**, identifiers as listed in the T1TAdb; **molecule_type**, localization of the hits in plasmid, chromosome, phage or prophage; **predicted_peptide**, amino-acid sequence of the predicted CDS; **nucleotide_locus**, coding strand DNA sequences ranging from 50 nt upstream to 200 nt downstream relative to the hit coordinates; **frame**, peptide coding frame relative to the nucleotide locus (only top frames); **strand**, peptide coding strand; **sd_start**, **sd_end**, **start_codon_start** and **stop_codon_end** indicate coordinates relative to the nucleotide locus marking the predicted Shine-Dalgarno (SD), start and stop codons respectively; **sd_sequence**, predicted SD, **peptide_start** and **peptide_end**, coordinates of the ORF relative to the accession (only for hits identified in the Refseq database); **peptide_prediction**, method for annotating the peptide in searches using nucleotide sequences; **Family** and **Genus**, classification according to the GTDB; **GenBank-Acc**, replicon GenBank accession; **AssAcc**, RefSeq assembly accession; **SequenceLength**, number of nt in the replicon; **refseq_chromosome**, replicon RefSeq accession of the chromosome in each assembly; columns starting with **NCBI** are Hok-related RefSeq annotations overlapping the predicted ORF; **predicted_orf**, DNA sequence corresponding to the predicted peptide; **loci_count_clus100**, number of loci containing each unique peptide sequence (clus100); **percent_in_mol_clus100**, percent of each unique sequence (clus100) by molecule type (rounded to 2 decimals); **unique_molecules_clus100**, list of molecule types where the clus100 is represented; **percent_per_genus_clus100**, percent of each unique sequence (clus100) per genus (rounded to 3 decimals); **rank_clus100_clus60**, ranking from most (0) to least (maximum number per clus100) represented sequence per clus60; **experimentally_tested**, sequences chosen for toxicity assays are marked with an X.

**Table S2 (separate file)**

**All experimentally tested Hok sequences.**

The columns are: **name**, code of the cloned plasmid; **cluster 60% _cluster 100%**, 60% and 100% identity cluster identifier to which each predicted peptide belongs; **cluster abundance**, LAC: low abundance cluster (only one sequence) and HAC: high abundance cluster (with more than 10 sequences); **predicted SD**, indicates the presence and sequence of predicted SD; **predicted orf**, DNA sequence corresponding to the predicted peptide; **predicted_peptide**, amino-acid sequence of the predicted CDS.

### Table S3. Strains

| Description | Reference |
|---|---|
| One Shot™ TOP10 Chemically Competent *E. coli* | Invitrogen |
| *Escherichia coli* BW25113 [pJAT13*araE*] $Erm^R$ | BW25113: DSMZ collection: DSM 27469, pJAT13*araE*: addgene item #18987 (1) |
| MG1655 | Lab collection |
| LR27: XTL632 *tetA-sacB*::*hok*_G202A/ΔSok | (2) |

### Table S4. Plasmids

| Name | Description | Resistance | Reference |
|---|---|---|---|
| pAZ3 | Empty vector with pBAD inducible promoter | $Cm^R$ | F. Darfeuille laboratory |
| LRP38 | pAZ3ΩTA05999<br>*hok*/Sok homolog TA05999 full loci cloned into pAZ3<br>https://d-lab.arna.cnrs.fr/display/display_feature_table/TA05999 | $Cm^R$ | This study |
| **Name** | **Cluster 60% _cluster 100%** | **Resistance** | **Reference** |
| LRP126 | pAZ3Ω47_842 | $Cm^R$ | This study |
| LRP127 | pAZ3Ω46_1073 | $Cm^R$ | This study |
| LRP131 | pAZ3Ω47_385_Flm | $Cm^R$ | This study |
| LRP132 | pAZ3Ω47_254 | $Cm^R$ | This study |
| LRP133 | pAZ3Ω35_697_HokD | $Cm^R$ | This study |
| LRP135 | pAZ3Ω93_1386 | $Cm^R$ | This study |
| LRP137 | pAZ3Ω72_1621 | $Cm^R$ | This study |
| LRP138 | pAZ3Ω38_529_HokC | $Cm^R$ | This study |
| LRP139 | pAZ3Ω15_1427 | $Cm^R$ | This study |
| LRP152 | pAZ3Ω78_1774 | $Cm^R$ | This study |
| LRP153 | pAZ3Ω47_235 | $Cm^R$ | This study |
| LRP154 | pAZ3Ω93_1170 | $Cm^R$ | This study |
| LRP155 | pAZ3Ω35_700 | $Cm^R$ | This study |
| LRP156 | pAZ3Ω93_1348 | $Cm^R$ | This study |
| LRP157 | pAZ3Ω72_1589 | $Cm^R$ | This study |
| LRP158 | pAZ3Ω27_135 | $Cm^R$ | This study |
| LRP160 | pAZ3Ω54_1205 | $Cm^R$ | This study |
| LRP161 | pAZ3Ω60_1235 | $Cm^R$ | This study |
| LRP162 | pAZ3Ω76_1683 | $Cm^R$ | This study |
| LRP163 | pAZ3Ω79_1563 | $Cm^R$ | This study |
| LRP164 | pAZ3Ω80_1515 | $Cm^R$ | This study |
| LRP165 | pAZ3Ω82_1289 | $Cm^R$ | This study |
| LRP166 | pAZ3Ω15_1414_PndA_R483 | $Cm^R$ | This study |
| LRP167 | pAZ3Ω93_1378_HokA | $Cm^R$ | This study |
| LRP168 | pAZ3Ω46_966_HokE | $Cm^R$ | This study |
| LRP169 | pAZ3Ω60_1482 | $Cm^R$ | This study |
| LRP170 | pAZ3Ω35_712 | $Cm^R$ | This study |
| LRP171 | pAZ3Ω46_982 | $Cm^R$ | This study |
| LRP172 | pAZ3Ω47_377_Hok_R1 | $Cm^R$ | This study |
| LRP183 | pAZ3Ω47_188_HokB | $Cm^R$ | This study |
| LRP184 | pAZ3Ω47_377_Hok_R1*Stop( | $Cm^R$ | This study |
| LRP185 | pAZ3Ω79_1555 | $Cm^R$ | This study |
| LRP204 | pAZ3Ω14_128 | $Cm^R$ | This study |
| LRP205 | pAZ3Ω21_260 | $Cm^R$ | This study |
| LRP206 | pAZ3Ω28_511 | $Cm^R$ | This study |
| LRP207 | pAZ3Ω32_557 | $Cm^R$ | This study |
| LRP208 | pAZ3Ω37_816 | $Cm^R$ | This study |
| LRP209 | pAZ3Ω41_889 | $Cm^R$ | This study |

| | | | |
|---|---|---|---|
| LRP210 | pAZ3Ω42_894 | $Cm^R$ | This study |
| LRP211 | pAZ3Ω45_907 | $Cm^R$ | This study |
| LRP212 | pAZ3Ω64_1457 | $Cm^R$ | This study |
| LRP213 | pAZ3Ω68_1529 | $Cm^R$ | This study |
| LRP214 | pAZ3Ω69_1540 | $Cm^R$ | This study |
| LRP215 | pAZ3Ω73_1644 | $Cm^R$ | This study |
| LRP216 | pAZ3Ω19_179 | $Cm^R$ | This study |
| LRP217 | pAZ3Ω47_377_Hok_R1*C31A | $Cm^R$ | This study |
| LRP218 | pAZ3Ω47_377_Hok_R1*E41A | $Cm^R$ | This study |
| LRP221 | pAZ3Ω47_377_Hok_R1*C16A | $Cm^R$ | This study |
| LRP222 | pAZ3Ω47_377_Hok_R1*L30A | $Cm^R$ | This study |
| LRP223 | pAZ3Ω47_377_Hok_R1*E32A | $Cm^R$ | This study |
| LRP223 | pAZ3Ω47_377_Hok_R1*T18A | $Cm^R$ | This study |
| LRP225 | pAZ3Ω66_1506 | $Cm^R$ | This study |
| LRP226 | pAZ3Ω47_377_Hok_R1*E49A | $Cm^R$ | This study |

### Table S5. Oligonucleotides

| Name | Sequence | Description |
|---|---|---|
| LR03 | aaatcaccagcaaacaccga | Primers used to amplify and clone the TA05999 *hok* mRNA in pAZ3 using EcoRI and HindIII restriction sites |
| LRO228 | ggctgtaagcttcaacagcaatgcttacgcataa | |
| LRO491 | aaccatagcgaaaatagtggcgcagtgtaattactgcttacactgtaagaac | Primers to amplify the LRP38 plasmid downstream the pBAD promoter and adding a strong RBS (taaggaggt)(3) |
| LRO608 | ttacactgcgccactattttcgctatggttatgcgtaagcattgctgttgaagcttgg ctgttttggcg | |
| | **Couples of primers to amplify *hok* homolog CDS (from Start to Stop codon) with homologous regions to LRP38 for AQUA cloning** | **clus60_clus100** |
| LRO465 | tgaattcacccgtttttaaggaggtaaaaaatgaaacgcaaccctctggtg | 47_842 |
| LRO580 | aaccatagcgaaaatagtggcgcagtgtaattacctggacgtgcaggccatg | |
| LRO469 | tgaattcacccgtttttaaggaggtaaaaaatgccgcagaaatatagattactttctt taatag | 93_1386 |
| LRO588 | aaccatagcgaaaatagtgcgcagtgtaattactcttttaaattgcaggctaaaaatg | |
| LRO483 | tgaattcacccgtttttaaggaggtaaaaaatgaaactgccgaaccaacc | 47_254 |
| LRO601 | aaccatagcgaaaatagtggcgcagtgtaactacttaccggattcgtagtc | |
| LRO589 | tgaattcacccgtttttaaggaggtaaaaaatgccacagcgaacgtttttaatg | 15_1427 |
| LRO590 | aaccatagcgaaaatagtggcgcagtgtaattaacgtttaacttcgtaggctaac | |
| LRO463 | tgaattcacccgtttttaaggaggtaaaaaatgaaactaccacgcagctctc | 47_385 Flm |
| LRO583 | aaccatagcgaaaatagtggcgcagtgtaactacttaccggattcgtaagc | |
| LRO475 | tgaattcacccgtttttaaggaggtaaaaaatgaaactaccgggaaacgcc | 47_235 |
| LRO593 | aaccatagcgaaaatagtggcgcagtgtaactacttaccgccggattcgtaag | |
| LRO478 | tgaattcacccgtttttaaggaggtaaaaaatgccaaaacgtactctgctg | 93_1170 |
| LRO598 | aaccatagcgaaaatagtggcgcagtgtaactaacgttttgcttcgtaagc | |
| LRO464 | tgaattcacccgtttttaaggaggtaaaaaatgctgacgaaatatgcccttg | 46_1073 |
| LRO581 | aaccatagcgaaaatagtggcgcagtgtaactacttcttcggttcgtaagc | |
| LRO467 | tgaattcacccgtttttaaggaggtaaaaaatgaagcagcataaggcgatg | 38_529 HokC |
| LRO586 | aaccatagcgaaaatagtggcgcagtgtaattactcggattcgtaagccg | |
| LRO474 | tgaattcacccgtttttaaggaggtaaaaaatgaagcagcaaaaggcgatg | 35_697 HokD |
| LRO592 | aaccatagcgaaaatagtggcgcagtgtaattactcctcaggttcgtaagc | |
| LRO477 | tgaattcacccgtttttaaggaggtaaaaaatgaagcagcaaaaggcgatg | 35_700 |
| LRO597 | aaccatagcgaaaatagtggcgcagtgtaattactcctcagattcgtaagctg | |
| LRO484 | tgaattcacccgtttttaaggaggtaaaaaatgaagcagcaaaaggcgatg | 35_712 |
| LRO600 | aaccatagcgaaaatagtggcgcagtgtaattacttctcagattcgtagtctacg | |
| LRO485 | tgaattcacccgtttttaaggaggtaaaaaatgtcgcaaaaatcgcttatcacc | 72_1589 |

| | | |
|---|---|---|
| LRO602 | aaccatagcgaaaatagtggcgcagtgtaactactgcttacactgtaagaac | |
| LRO594 | tgaattcacccgtttttaaggaggtaaaaaatgtcgcaaaaatcgcttac | 72_1621 |
| LRO595 | aaccatagcgaaaatagtggcgcagtgtaattactgcttacactgtaagaac | |
| LRO473 | tgaattcacccgtttttaaggaggtaaaaaatgacgccattaaaaactgcgttag | 78_1774 |
| LRO591 | aaccatagcgaaaatagtggcgcagtgtaattagcctgcgttgcaggctaattttg | |
| LRO476 | tgaattcacccgtttttaaggaggtaaaaaatgccgcaaaagtatctgttgtttg | 93_1348 |
| LRO596 | aaccatagcgaaaatagtggcgcagtgtaattactgtttaatatcacaggctaag | |
| LRO480 | tgaattcacccgtttttaaggaggtaaaaaatgcagcaaaaacgggtcg | 60_1482 |
| LRO599 | aaccatagcgaaaatagtggcgcagtgtaattaccgttcggattcgtaag | |
| LRO468 | tgaattcacccgtttttaaggaggtaaaaaatgctgacaaaatatgcccttgtg | 46_982 |
| LRO587 | aaccatagcgaaaatagtggcgcagtgtaactacttcttcgattcgtaagc | |
| LRO606 | tgaattcacccgtttttaaggaggtaaaaaatgaagcacaaccctctggtg | 47_188 HokB |
| LRO607 | ccatagcgaaaatagtggcgcagtgtaattacctggacgtgcaggccatg | |
| LRO609 | tgaattcacccgtttttaaggaggtaaaaaaggagccacacaaaccag | 27_135 |
| LRO610 | aaccatagcgaaaatagtggcgcagtgtaatactcggatccgtaggccatg | |
| LRO615 | tgaattcacccgtttttaaggaggtaaaaaatgccaaaaagtaaaaccgc | 54_1205 |
| LRO616 | aaccatagcgaaaatagtggcgcagtgtaactacttcatacgtgtttcgtag | |
| LRO617 | tgaattcacccgtttttaaggaggtaaaaaatgccgaacaaacggagcctg | 60_1235 |
| LRO618 | aaccatagcgaaaatagtgggccagtgtaattaccgttcggattcgtaagc | |
| LRO619 | tgaattcacccgtttttaaggaggtaaaaaatgactaagcttgcccttattg | 76_1683 |
| LRO620 | aaccatagcgaaaatagtggcgcagtgtaactatttatcgcaagataaattc | |
| LRO621 | tgaattcacccgtttttaaggaggtaaaaaatgtcgcaaaaaccgttaaaaac | 79_1563 |
| LRO622 | aaccatagcgaaaatagtggcgcagtgtaattaccgcttacactgtaagaac | |
| LRO623 | tgaattcacccgtttttaaggaggtaaaaaatgcgcttaaggcatgtgtttc | 80_1515 |
| LRO624 | aaccatagcgaaaatagtggcgcagtgtaattaccgcttcacctcacacgc | |
| LRO625 | tgaattcacccgtttttaaggaggtaaaaaatgccgcaaaaaacgatcattg | 82_1289 |
| LRO626 | aaccatagcgaaaatagtggcgcagtgtaattacggtttacactgcaagaa | |
| LRO627 | tgaattcacccgtttttaaggaggtaaaaaatgccacagcgaacgtttttaatg | 15_1414 PndA from plasmid R483 |
| LRO628 | aaccatagcgaaaatagtggcgcagtgtaattaacgtttaacttcgtaggc | |
| LRO639 | tgaattcacccgtttttaaggaggtaaaaaatgccgcagaaatatagattac | 93_1378 HokA |
| LRO640 | aaccatagcgaaaatagtggcgcagtgtaattactctttcaatttgcagg | |
| LRO641 | tgaattcacccgtttttaaggaggtaaaaaatgctgacgaaatatgcccttg | 46_966 HokE |
| LRO642 | aaccatagcgaaaatagtggcgcagtgtaactacttcttcggttcgtaagc | |
| LRO579 | aaccatagcgaaaatagtggcgcagtgtaactacttaccggattcgtaagc | 47_377 Hok from plasmid R1 and R1*stop |
| LRO461 | tgaattcacccgtttttaaggaggtaaaaaatgaaactaccacgaagttccc | |
| LRO584 | tgaattcacccgtttttaaggaggtaaaaaatgtcgcaaaaatcgctatc | 79_1555 |
| LRO585 | aaccatagcgaaaatagtggcgcagtgtaattactgcttacactgtaagaac | |
| LRO643 | tatcgcaactctctactgtttg | To amplify CDS of Hok mutants or homologs from low abundance clusters. The same sequence was added upstream and downstream the CDS to amplify all sequences with the same oligos |
| LRO644 | caaaacagccaagcttcaacag | |
| **Name** | **Sequence** | **Description** |
| LRO273 | cgtcacactttgctatgcca | Primers used for colony PCR and sequencing |
| LRO497 | ctgaaaatcttctctcatccg | |

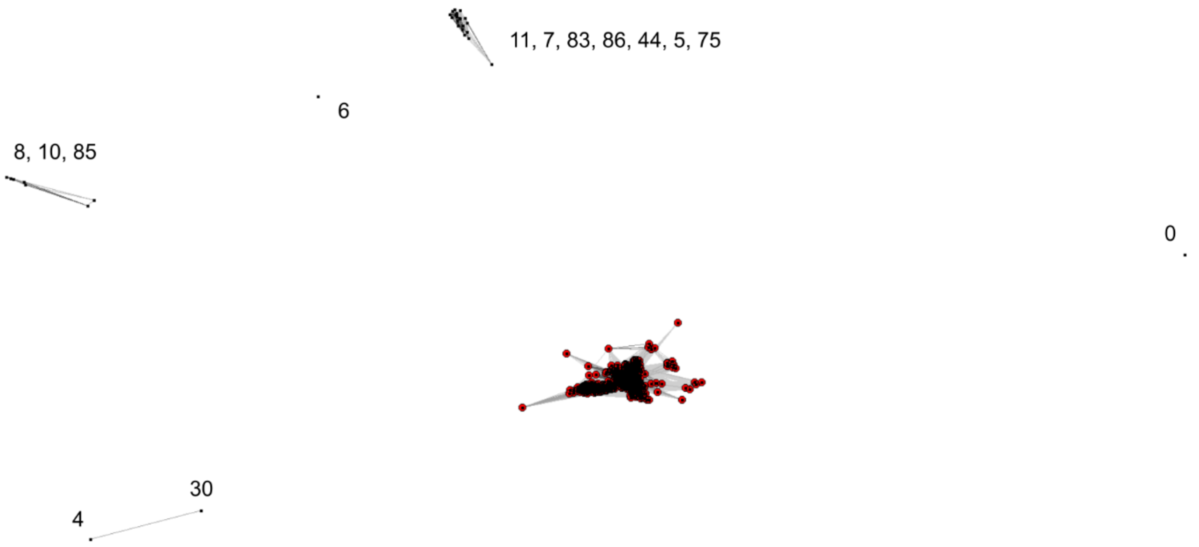


**Figure S1. Hok outlier removal by clustering.**

Graphical representation of sequence similarities visualized using CLANS. Protein sequences were compared in an all-against-all BLAST search and the resulting E-values were used to calculate pairwise attractive forces between sequences. Lower E-values correspond to stronger attractive forces and smaller distances. Each node in the graph represents a distinct peptide sequence. Clusters indicate groups of sequences with higher similarity. Edges are drawn for pairwise comparisons with a p-value lower than $1x10^{-4}$. Red dots mark the selected sequences. Numbers correspond the 60% identity cluster of the outliers, which are not connected with the main cluster and were removed from the analysis.

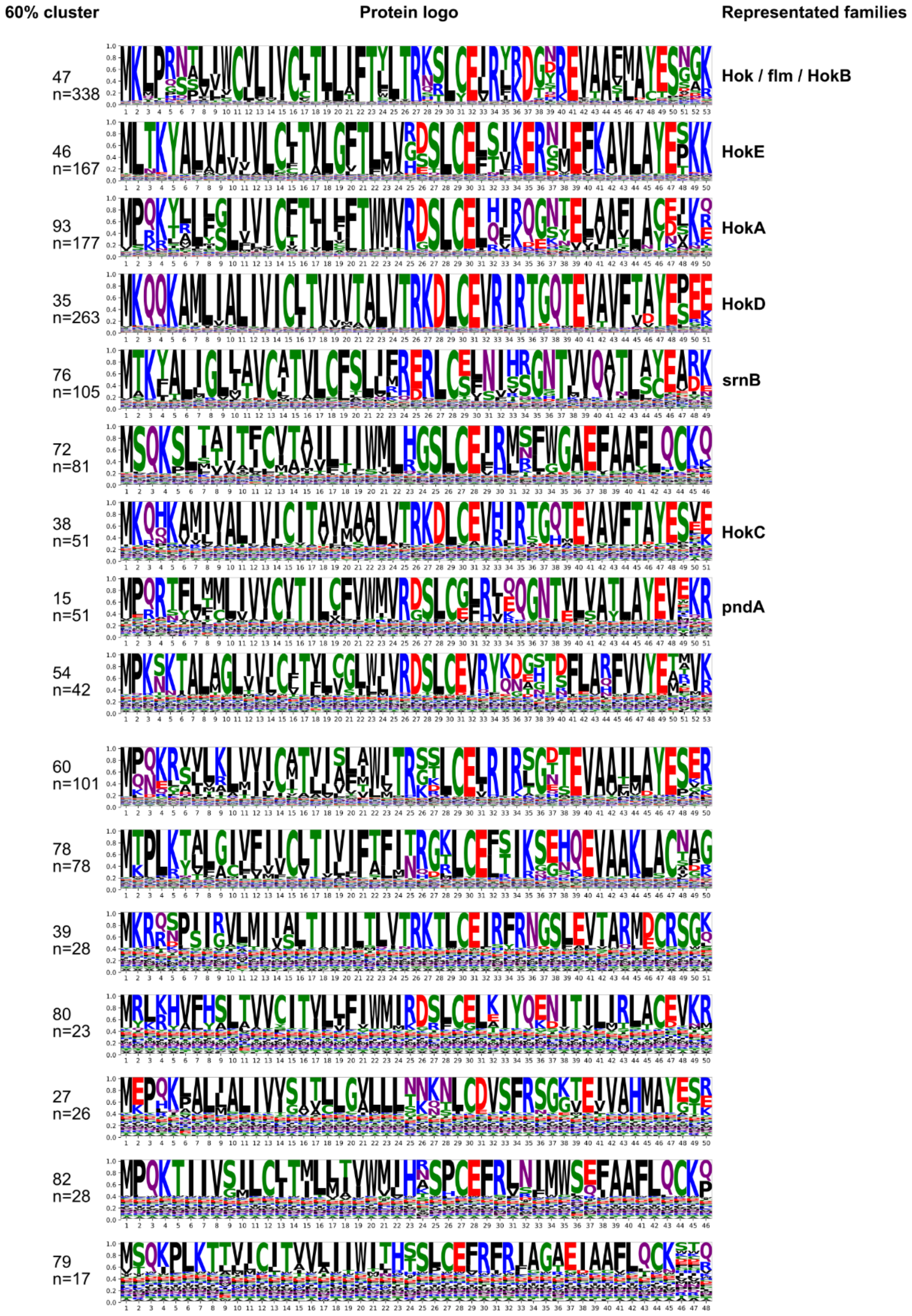


**Figure S2. Amino acid conservation analysis per cluster.**

Sequence logos representing Hok alignments for each 60% identity cluster with more than 10 unique sequences. The height of each letter at a given position reflects its relative frequency. The number of the 60% identity cluster is indicated. Logos display conserved positions with less than 90% gaps across aligned sequences. The number of distinct sequences included in the alignment is indicated (n).

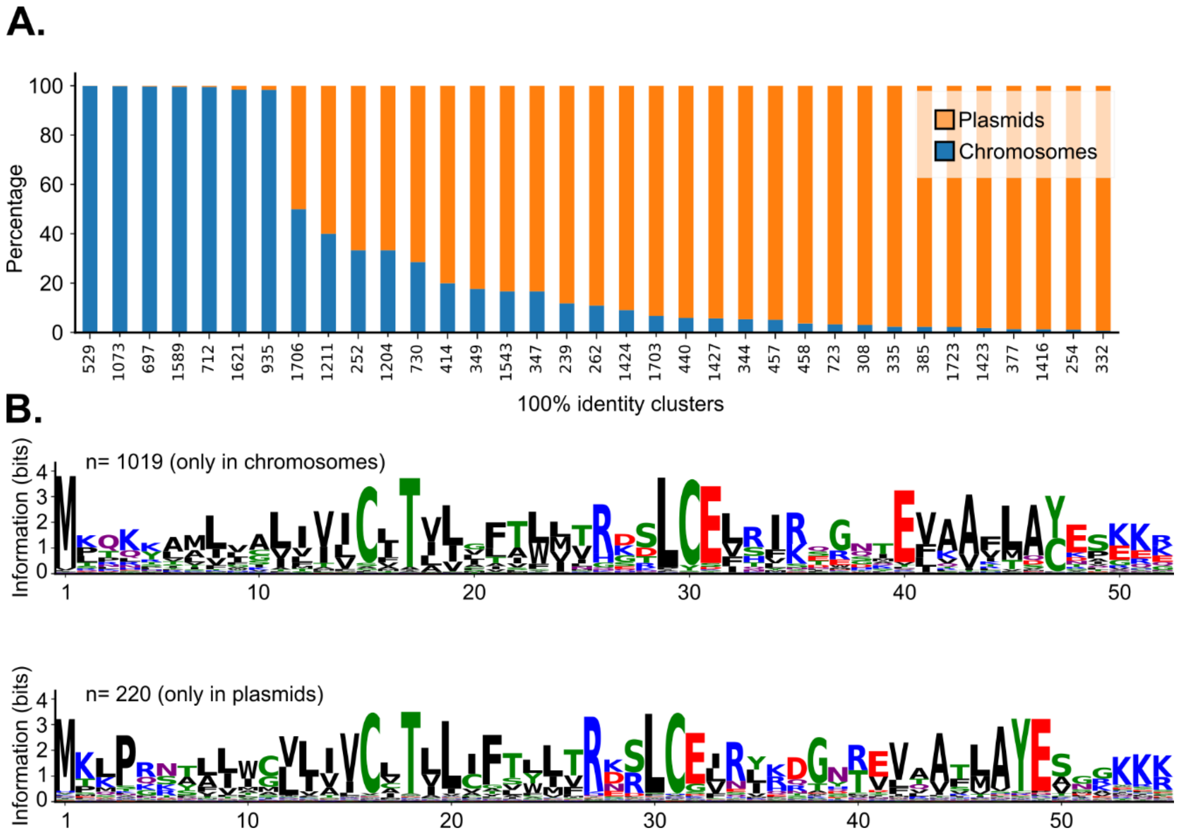


**Figure S3. Comparison of Hok sequences found in plasmids and chromosomes.**

**A.** Bar plot displaying the percentage of loci containing each Hok unique sequence in plasmids and chromosomes. The x-axis shows the identifier of each unique sequence (100% identity clusters). Only sequences present in both, plasmids and chromosomes, are represented. **B.** Sequence logos from protein alignments of homologs found only in plasmids (top) or chromosomes (bottom). The height of each letter at a given position reflects its relative frequency, while the overall stack height indicates the information content at that position (measured in bits). Logos display conserved positions with less than 90% gaps across aligned sequences. The number of distinct sequences included in the alignment is indicated (n).

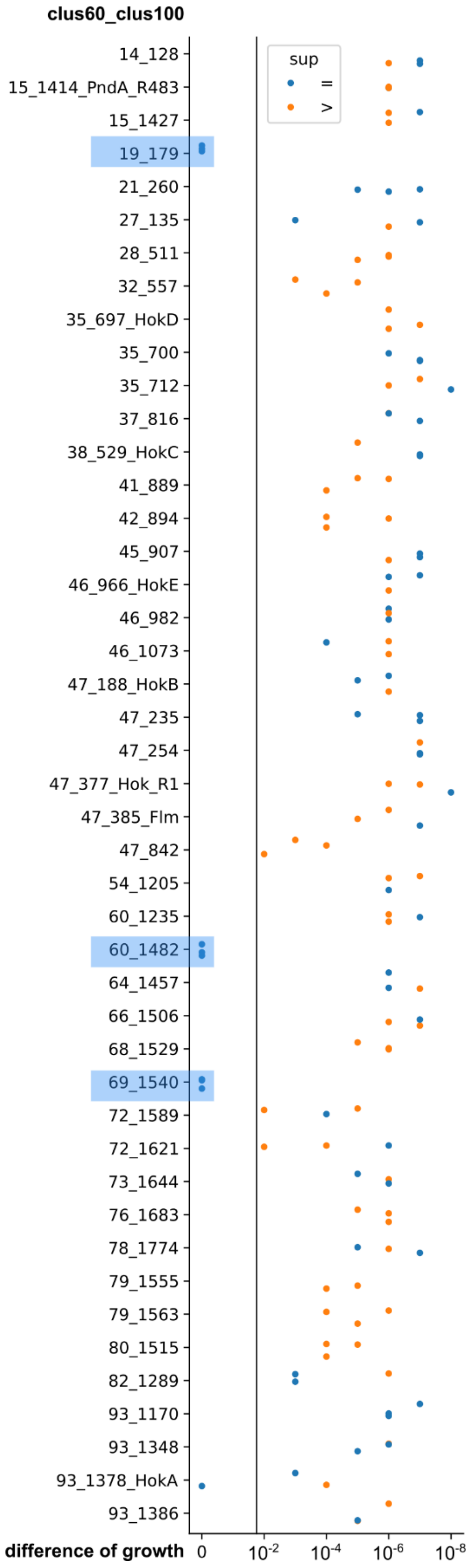


**Figure S4. Comparison of Hok homolog toxicity.**
Stripplot showing the toxicity of each Hok peptide homolog. Toxicity was measured as the difference ($log_{10}$ scale) between the highest dilution showing growth under inducing conditions (arabinose) and the highest dilution showing growth under repressing conditions (glucose). Blue dots indicate exact differences, while orange dots indicate that the growth difference exceeded the measurable range. Only three homologs, highlighted in blue, did not exhibit toxicity.

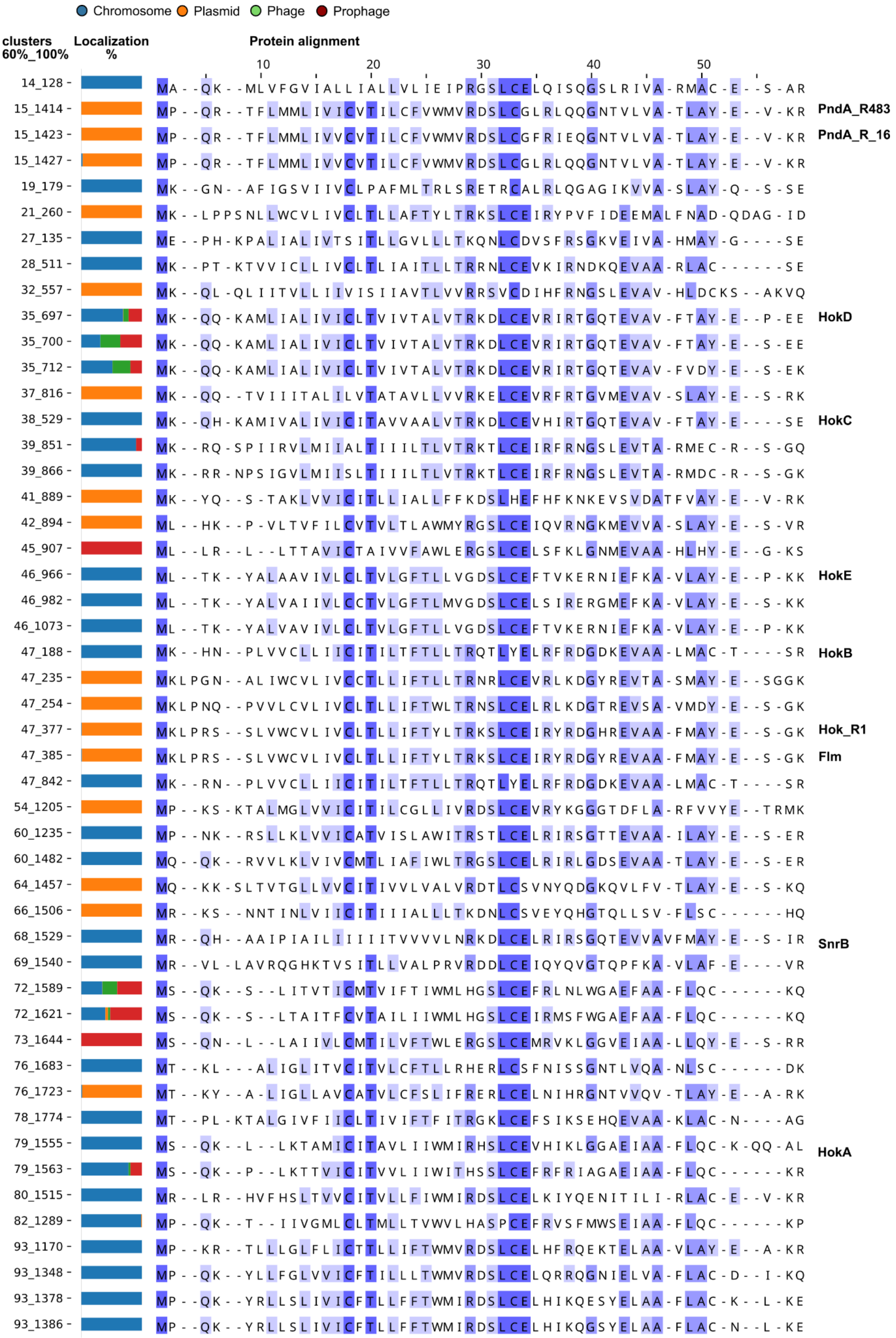
Chromosome Plasmid Phage Prophage
clusters 60%_100%
Localization %
Protein alignment
10 20 30 40 50
14_128 MA--QK--MLVFGVIALLIALLVLIEIPRGSLCELQISQGSLRIVA-RMAC-E--S-AR
15_1414 MP--QR--TFLMMLIVICVTILCFVWMVRDSLCGLRLQQGNTVLVA-TLAY-E--V-KR PndA_R483
15_1423 MP--QR--TFLMMLIVVCVTILCFVWMVRDSLCGFRIEQGNTVLVA-TLAY-E--V-KR PndA_R_16
15_1427 MP--QR--TFLMMLIVVCVTILCFVWMVRDSLCGLRLQQGNTVLVA-TLAY-E--V-KR
19_179 MK--GN--AFIGSVIIVCLPAFMLTRLSRETRCALRLQGAGIKVVA-SLAY-Q--S-SE
21_260 MK--LPPSNLLWCVLIVCLTLLAFTYLTRKSLCEIRYPVFIDEEMALFNAD-QDAG-ID
27_135 ME--PH-KPALIALIVTSITLLGVLLLTKQNLCDVSFRSGKVEIVA-HMAY-G----SE
28_511 MK--PT-KTVVICLLIVCLTLIAITLLTRRNLCEVKIRNDKQEVAA-RLAC------SE
32_557 MK--QL-QLIITVLLIIVISIIAVTLVVRRSVCDIHFRNGSLEVAV-HLDCKS--AKVQ
35_697 MK--QQ-KAMLIALIVICLTVIVTALVTRKDLCEVRIRTGQTEVAV-FTAY-E--P-EE HokD
35_700 MK--QQ-KAMLIALIVICLTVIVTALVTRKDLCEVRIRTGQTEVAV-FTAY-E--S-EE
35_712 MK--QQ-KAMLIALIVICLTVIVTALVTRKDLCEVRIRTGQTEVAV-FVDY-E--S-EK
37_816 MK--QQ--TVIIITALILVTATAVLLVVRKELCEVRFRTGVMEVAV-SLAY-E--S-RK
38_529 MK--QH-KAMIVALIVICITAVVAALVTRKDLCEVHIRTGQTEVAV-FTAY-E----SE HokC
39_851 MK--RQ-SPIIRVLMIIALTIIILTLVTRKTLCEIRFRNGSLEVTA-RMEC-R--S-GQ
39_866 MK--RR-NPSIGVLMIISLTIIILTLVTRKTLCEIRFRNGSLEVTA-RMDC-R--S-GK
41_889 MK--YQ--S-TAKLVVICITLLIALLFFKDSLHEFHFKNKEVSVDATFVAY-E--V-RK
42_894 ML--HK--P-VLTVFILCVTVLTLAWMYRGSLCEIQVRNGKMEVVA-SLAY-E--S-VR
45_907 ML--LR--L--LTTAVICTAIVVFAWLERGSLCELSFKLGNMEVAA-HLHY-E--G-KS
46_966 ML--TK--YALAAVIVLCLTVLGFTLLVGDSLCEFTVKERNIEFKA-VLAY-E--P-KK HokE
46_982 ML--TK--YALVAIIVLCCTVLGFTLMVGDSLCELSIRERGMEFKA-VLAY-E--S-KK
46_1073 ML--TK--YALVAVIVLCLTVLGFTLLVGDSLCEFTVKERNIEFKA-VLAY-E--P-KK
47_188 MK--HN--PLVVCLLIICITILTFTLLTRQTLYELRFRDGDKEVAA-LMAC-T----SR HokB
47_235 MKLPGN--ALIWCVLIVCCTLLIFTLLTRNRLCEVRLKDGYREVTA-SMAY-E--SGGK
47_254 MKLPNQ--PVVLCVLIVCLTLLIFTWLTRNSLCELRLKDGTREVSA-VMDY-E--S-GK
47_377 MKLPRS--SLVWCVLIVCLTLLIFTYLTRKSLCEIRYRDGHREVAA-FMAY-E--S-GK Hok_R1
47_385 MKLPRS--SLVWCVLIVCLTLLIFTYLTRKSLCEIRYRDGYREVAA-FMAY-E--S-GK Flm
47_842 MK--RN--PLVVCLLIICITILTFTLLTRQTLYELRFRDGDKEVAA-LMAC-T----SR
54_1205 MP--KS-KTALMGLVVICITILCGLLIVRDSLCEVRYKGGGTDFLA-RFVVYE--TRMK
60_1235 MP--NK--RSLLKLVVICATVISLAWITRSTLCELRIRSGTTEVAA-ILAY-E--S-ER
60_1482 MQ--QK--RVVLKLVIVCMTLIAFIWLTRGSLCELRIRLGDSEVAA-TLAY-E--S-ER
64_1457 MQ--KK-SLTVTGLLVVCITIVVLVALVRDTLCSVNYQDGKQVLFV-TLAY-E--S-KQ
66_1506 MR--KS--NNTINLVIICITIIIALLLTKDNLCSVEYQHGTQLLSV-FLSC------HQ
68_1529 MR--QH--AAIPIAILIIIIITVVVVLNRKDLCELRIRSGQTEVVAVFMAY-E--S-IR SnrB
69_1540 MR--VL-LAVRQGHKTVSITLLVALPRVRDDLCEIQYQVGTQPFKA-VLAF-E----VR
72_1589 MS--QK--S--LITVTICMTVIFTIWMLHGSLCEFRLNLWGAEFAA-FLQC------KQ
72_1621 MS--QK--S--LTAITFCVTAILIIWMLHGSLCEIRMSFWGAEFAA-FLQC------KQ
73_1644 MS--QN--L--LAIIVLCMTILVFTWLERGSLCEMRVKLGGVEIAA-LLQY-E--S-RR
76_1683 MT--KL---ALIGLITVCITVLCFTLLRHERLCSFNISSGNTLVQA-NLSC------DK
76_1723 MT--KY--A-LIGLLAVCATVLCFSLIFRERLCELNIHRGNTVVQV-TLAY-E--A-RK
78_1774 MT--PL-KTALGIVFIICLTIVIFTFITRGKLCEFSIKSEHQEVAA-KLAC-N----AG
79_1555 MS--QK--L--LKTAMICITAVLIIWMIRHSLCEVHIKLGGAEIAA-FLQC-K-QQ-AL HokA
79_1563 MS--QK--P--LKTTVICITVVLIIWITHSSLCEFRFRIAGAEIAA-FLQC------KR
80_1515 MR--LR--HVFHSLTVVCITVLLFIWMIRDSLCELKIYQENITILI-RLAC-E--V-KR
82_1289 MP--QK--T--IIVGMLCLTMLLTVWVLHASPCEFRVSFMWSEIAA-FLQC------KP
93_1170 MP--KR-TLLLGLFLICTTLLIFTWMVRDSLCELHFRQEKTELAA-VLAY-E--A-KR
93_1348 MP--QK--YLLFGLVVICFTILLLTWMVRDSLCELQRRQGNIELVA-FLAC-D--I-KQ
93_1378 MP--QK--YRLLSLIVICFTLLFFTWMIRDSLCELHIKQESYELAA-FLAC-K--L-KE
93_1386 MP--QK--YRLLSLIVICFTLLFFTWMIRDSLCELHIKQGSYELAA-FLAC-N--L-KE

**Figure S5. Alignment of Hok homologs selected for toxicity assays.**
Protein alignments using Muscle5 of selected and precedently characterized Hok homologs visualized using Jalview, colored by percentage identify. One or several representative sequences of each cluster were tested experimentally except for clus39, for which we did not obtain unmutated plasmid constructions. The bar plot on the left represents the percentage of loci in each 60% identity cluster localized in chromosomes, plasmids, prophages or phages.

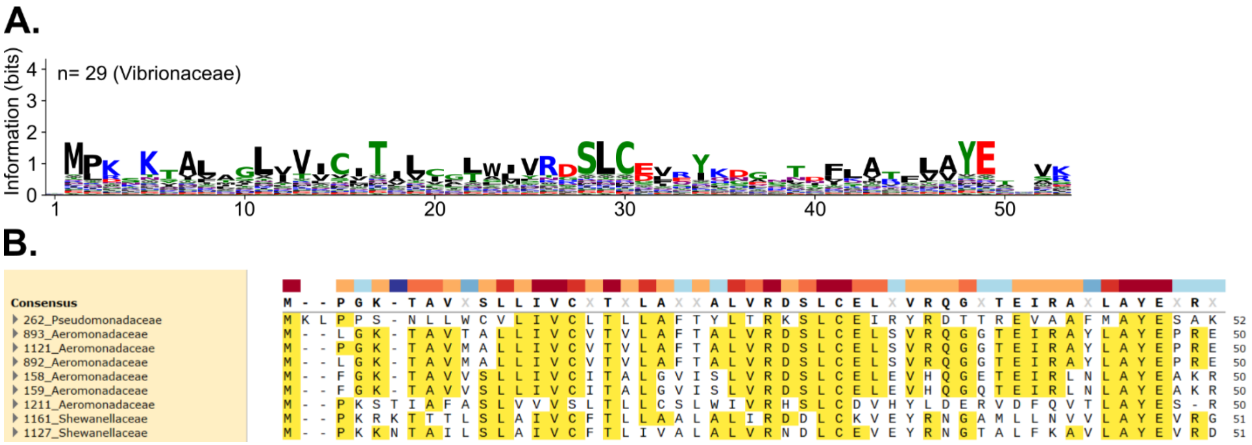


**Figure S6. Hok sequences present outside the Enterobacteriaceae family.**

**A.** Sequence logo from a protein alignment of the 29 homologs found in the Vibrionaceae family. The height of each letter at a given position reflects its relative frequency, while the overall stack height indicates the information content at that position (measured in bits). **B.** Peptides alignment of Hok homologs found in the Pseudomonadaceae, Aeromonadaceae and Shewanellaceae families, visualized using SnapGene. Only positions with less than 90% gaps across aligned sequences are shown in this figure.

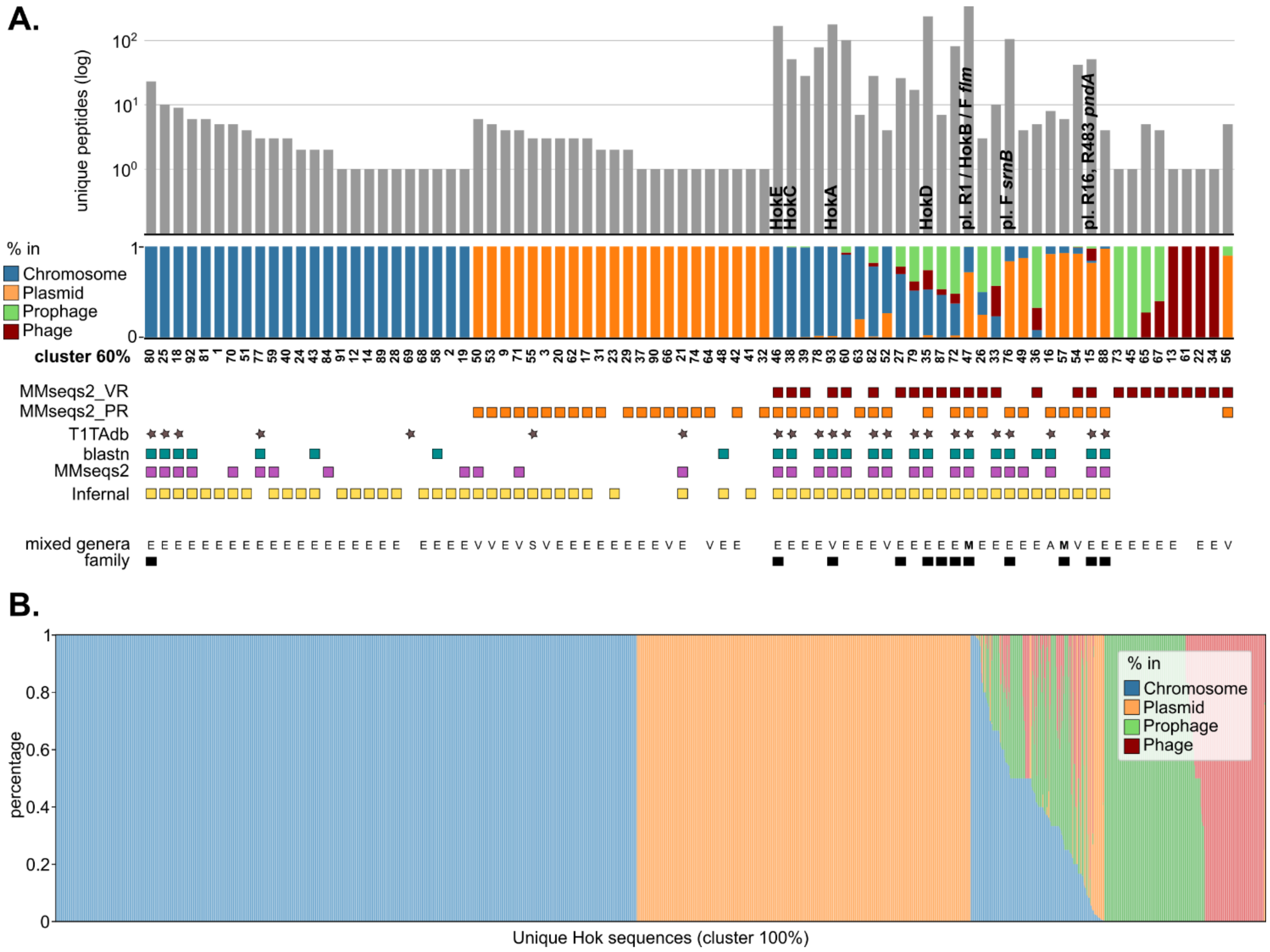


**Figure S7. Hok homolog identification by method in different locations.**

**A.** Information on the 60% identity clusters of Hok homologs. Clusters that contain sequences present in more than one genus are marked as "mixed genera" with a black square (bottom). The bacterial families represented in each cluster are marked: E (Enterobacteriaceae), V (Vibrionaceae), S (Shewanellaceae), A (Aeromonadaceae) and M (mixed, containing sequences from more than one family). Colored squares above the cluster numbers indicate the method they were identified by: MMseqs2 (protein sequence, purple), blastn (mRNA sequence, green) or infernal (mRNA structure and sequence, yellow) on complete bacterial genomes in the RefSeq database; MMseqs2 on plasmids in the IMG_PR database (orange) or phages in the IMG_VR database (brown); or present in the T1TAdb (gray star). Bars in middle show the percentage of loci in each cluster that are found in chromosomes (blue), plasmids (orange), prophages (green) or phages (dark red). The top bars indicate the number of unique peptides per cluster. The names of studied Hok homologs are written on top of the bar of their corresponding cluster. Note that clusters 25,70,52,2,23,21,20,9 and 41 were found with Infernal only when the structure was accounted for, but not when using sequence alone. There is a total of 80 clusters (79 identified here plus 1 cluster with a sequence from the T1TAdb not identified in our search).
**B.** Barplot showing the percentage of loci for each unique sequence (cluster 100%) that are found in chromosomes (blue), plasmids (orange), prophages (green) or phages (dark red).

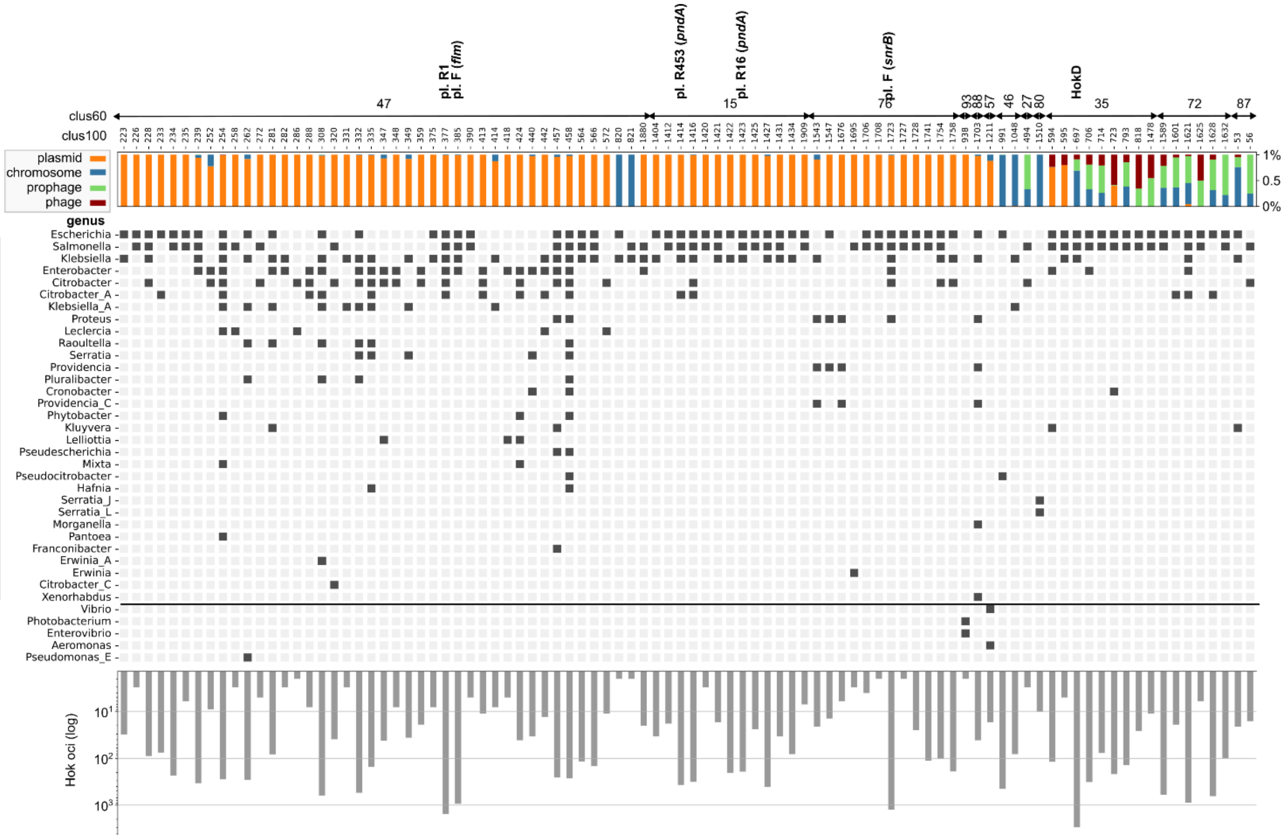


**Figure S8. Identical Hok sequences shared by different genera.**

Each column represents a distinct protein sequences with the unique sequence identifier (clus100) and their corresponding 60% identity cluster (clus60) indicated at the top. The bar plot on the top shows the percentage of loci for each identical sequence found in chromosomes (blue), plasmids (orange), prophages (green) or phages (dark red). The matrix in the middle indicates whether the sequence is present (dark square) or absent (light gray) in each genus from the Enterobacteriaceae family (top rows) or other families (5 last rows). The inverted bar plot at the bottom shows the number of loci encoding each sequence.

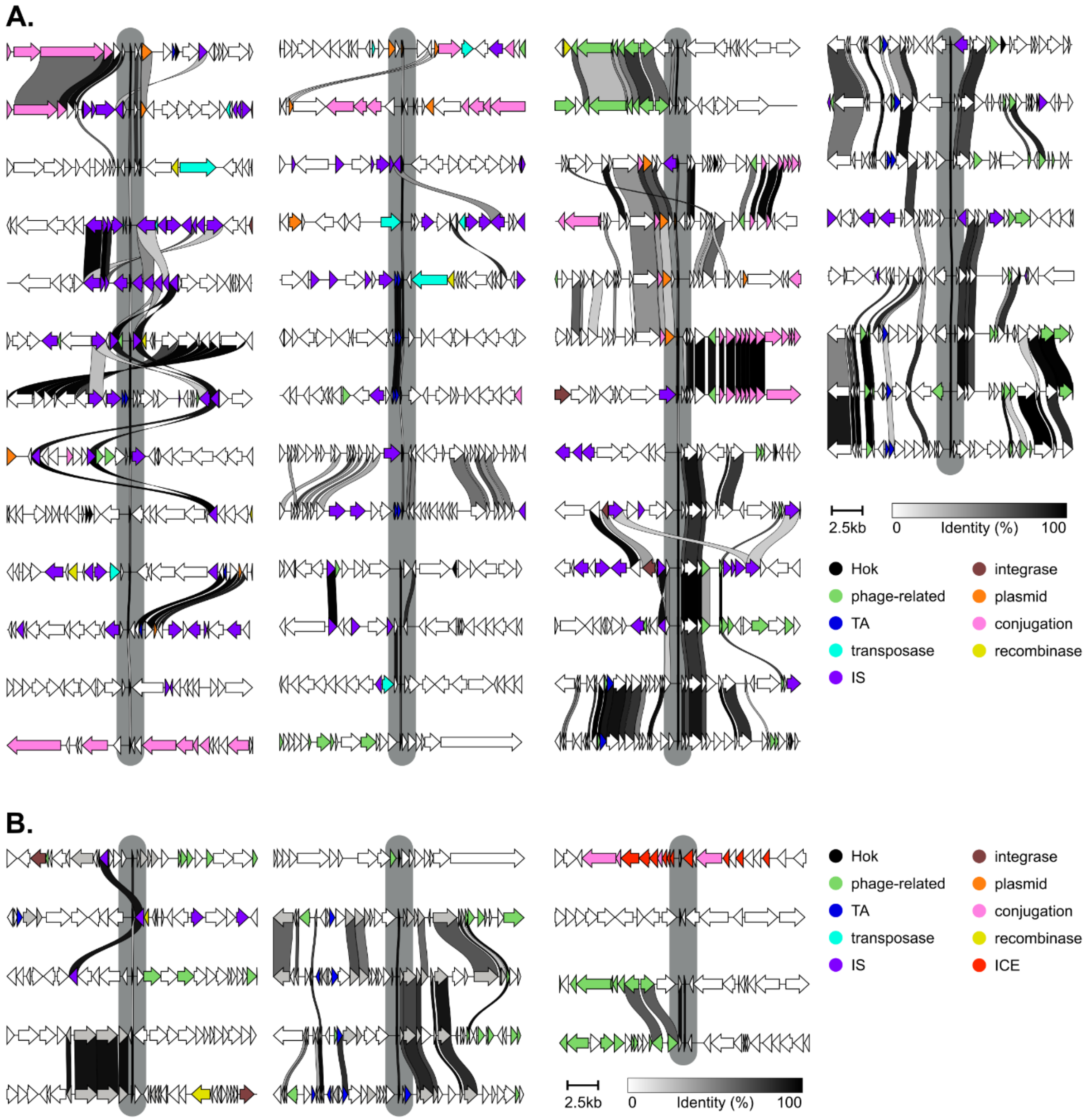


**Figure S9. Genomic context analysis of Hok sequences shared across genera.**
Functional annotation of the genes found in the 20 kb region adjacent to *hok* (in the middle of the gray highlight) in selected chromosomes. **A.** Loci encoding Hok sequences that were also found in plasmids. **B.** Loci encoding Hok sequences only found in chromosomes. Annotation and visualization were done using clinker. Genes with annotations related to mobile genetic elements were colored as indicated in the legend. Similarity between genes in adjacent loci are represented by the ribbons, with darker lines indicating higher amino-acid similarity (as shown in the heatmap bar). Genes are drawn to scale. Scale bar represents 2.5 kb.

**Supplementary references**


1. Khlebnikov,A., Datsenko,K.A., Skaug,T., Wanner,B.L. and Keasling,J.D. (2001) Homogeneous expression of the P(BAD) promoter in Escherichia coli by constitutive expression of the low-affinity high-capacity AraE transporter. *Microbiol. Read. Engl.*, **147**, 3241–3247.
2. Le Rhun,A., Tourasse,N.J., Bonabal,S., Iost,I., Boissier,F. and Darfeuille,F. (2023) Profiling the intragenic toxicity determinants of toxin-antitoxin systems: revisiting hok/Sok regulation. *Nucleic Acids Res.*, **51**, e4.
3. Chen,H., Bjerknes,M., Kumar,R. and Jay,E. (1994) Determination of the optimal aligned spacing between the Shine-Dalgarno sequence and the translation initiation codon of Escherichia coli mRNAs. *Nucleic Acids Res.*, **22**, 4953–4957.
